# Rapid Adaptation of Cellular Metabolic Rate to the MicroRNA Complements of Mammals and its Relevance to the Evolution of Endothermy

**DOI:** 10.1101/2022.11.24.517858

**Authors:** Bastian Fromm, Thomas Sorger

**Affiliations:** The Arctic University Museum of Norway, UiT- The Arctic University of Norway, 9037 Tromsø, Norway; Department of Biology, Roger Williams University, Bristol, Rhode Island 02809, USA

## Abstract

The metabolic efficiency of mammalian cells depends on attenuation of intrinsic translation noise by microRNAs. We devised a metric of cellular metabolic rate (*cMR)*, *rMR*/*M*^exp^ optimally fit to the number of microRNA families (*miRNA.Fam*), that is robust to variation in mass and sensitive to body temperature, consistent with the Heat Dissipation Limit Theory of Speakman and Król (2010)^1^. Using *miRNA.Fam* as predictor, an Ornstein-Uhlenbeck process of stabilizing selection, with an adaptive shift at the divergence of Boreoeutheria, accounted for 95% of the variation in *cMR* across mammals. Branchwise rates of evolution of *cMR*, *miRNA.Fam* and body temperature concurrently increased 6- to 7-fold at the divergence of Boreoeutheria, independent of mass. Cellular *MR* variation across placental mammals was also predicted by the sum of model conserved microRNA-target interactions, revealing an unexpected degree of integration of the microRNA-target apparatus into the energy economy of the mammalian cell.

## INTRODUCTION

Comparative phylogenetic models envision adaptive landscapes that can change gradually or abruptly^2, 3, 4, 5, 6, 7, 8, 9, 10, 11, 12^. The mechanism for traversing the landscape can be either random and clock-like, simulated by a Brownian motion process^13^, or deterministic and adaptive, such as the Ornstein-Uhlenbeck model of stabilizing selection^5, 8^. One candidate for an adaptive shift is the combination of permanent homeothermy, fast specific metabolic rates (*sMR*s), and mean body temperatures (*T*_b_’s) > 36°C among most orders of Euarchontoglires and those orders of Laurasiatheria that diverged more recently than Chiroptera. This energy regime contrasts with that found in Atlantogenata and earlier-branching orders of Boreoeutheria (Eulipotyphla, Chiroptera), with mean *T*_b_’s < 36°C, overlapping those of marsupials (Supplementary Figure S1), and that generally exhibit torpor, hibernation, and/or behavioral thermoregulation, reliant as they are on low-energy density insectivory or foraging^14, 15, 16, 17, 18, 19, 20, 21, 22^. The emergence of permanent homeothermy associated with fast *sMR*s, in birds as well as in mammals, can be understood as an endpoint in the evolution of the regulation of body temperature, the ultimate safeguarding of development from external perturbations, thereby allowing a more forward-looking and efficient allocation of energy among somatic maintenance, reproductive reserves, and parental care^23, 24, 25, 26^.

In large mammalian trees, the rate of evolution of basal *MR* (*bMR*) was more variable and better correlated with ambient temperature (*T*_a_) than was the rate of *T*_b_ evolution^27^, making it more likely to have been the target of selection. Overall *bMR* depends on mass, hence *sMR*, as well as on mass-independent factors, such as thermal conductance 26^28, 29^. Cellular *MR* accounts for 90% of *bMR*^30^ and depends on the rate of protein synthesis, which drives 20% of ATP turnover^31^. The rate of protein synthesis is, in turn, dictated by cell size, which is highly conserved in mammals^32, 33, 34^ (for reviews^35, 36^). Matching of the rate of protein synthesis to cell size is predicted by models in which the rate of translation in exponentially growing cells is limited by competition among mRNAs for ribosomes^37, 38^. A slow rate of translation relative to transcription minimizes energy losses due to both error correction by kinetic proofreading^39, 40, 41, 42, 43, 44, 45^, as well as to intrinsic translation noise^38, 44, 46^.

We propose that the fitness benefits of faster *sMR*s and higher *T*_b_’s would have been offset without compensatory adaptations to limit increases in energy dissipation due to the cost of kinetic proofreading and translation noise. The interactions between microRNAs and their messenger RNA targets constitute a well-established mechanism to attenuate intrinsic translation noise^47, 48^. The distribution of the rates of translation of low- and high-abundance mRNAs conform to the predicted outcome of global activity of microRNAs^49, 50^, which preferentially repress the translation of low-abundance transcripts^51^. Furthermore, these relative rates of translation appear to have evolved under an energy efficiency constraint^50^, which can be understood in light of the 10- to 100-fold higher thermodynamic cost per gene of translation relative to transcription in eukaryotes^52, 53^. Assuming conservatively that global microRNA activity reduces the global rate of translation by 25%^54, 55, 56, 57, 58, 59^, a 10-fold increase in cellular *MR*, without a corresponding increase in global microRNA activity, would incur a 6% loss of efficiency of cellular *MR* due to protein overproduction (5.4% of overall *rMR*), as well as add to the cost of protein degradation^60, 61^. While not directly comparable, it is worth noting that losses of fitness of up to 3% were found when the intrinsic translation noise of individual, constitutive proteins in yeast was doubled^62^.

Consistent with an adaptation to maintain energy efficiency, the complements of conserved microRNAs and density of conserved microRNA target sites in placental mammals are larger than those of metatheria, with marked increases among permanently homeothermic, late-branching Boreoeutheria^63, 64, 65, 66, 67^ (Supplementary Tables S1 and S2). We postulated that the emergence of permanent homeothermy by thermoregulation in fast *sMR* Boreoeutheria represented the appearance of a new optimum in the adaptive landscape, characterized by coordinated increases in microRNA repertoires, microRNA-target interactions, cellular *MR*s and *T*_b_. By assuming that global microRNA activity is proportional to the number of microRNA families (*miRNA.Fam*), we devised a new metric to compare cellular *MR* across species. Rather than inferring *sMR* from the slope of log *rMR versus* log *M*, we used variable allometric scaling to optimally fit *rMR*/*M*^exp^ to the distribution of *miRNA.Fam*. We interpret this *rMR*/*M*^opt^ as the global rate of microRNA-mediated repression of translation, which we assume is proportional to cellular metabolic rate and designate as *cMR*, in order to avoid confusion with conventional *sMR*. We demonstrate below that this metric of cellular *MR* has desirable properties compared to inference from the ratio log *rMR versus* log *M*, such as robustness to variation in mass and sensitivity to body temperature of the allometric exponent, hence the implied ratio of surface area (a measure of the of rate heat dissipation) to mass or volume (a measure of the rate of heat generation)^68, 69^. Importantly, the variation across species of *cMR* with respect to *miRNA.Fam* (or any other covariate) is homoscedastic (Supplementary Figure S2), and therefore compatible with regression models^70^, allowing us to test the following predictions:

i. The evolution of *cMR* with respect to *miRNA.Fam* will conform better to a deterministic (Ornstein-Uhlenbeck) model of adaptation than to a model of random drift by Brownian motion;
ii. The distribution of optimally-scaled *cMR* will be best simulated by an OU process with an adaptive shift corresponding to the divergence of Boreoeutheria;
iii. Since genetic correlations between activity, mass and *MR* have been attributed to selection^71, 72^, an adaptive shift to permanent homeothermy will be evidenced by increases of two-fold or more^73^ in the rates of evolution of *miRNA.Fam*, *cMR* and *T*_b_;
iv. The relative increase in *cMR* among placental mammals will correspond to approximately 20% of the reduction in translation cost that can be attributed to the sum of all model microRNA-target interactions;
v. The allometric scaling of *cMR* will depend more on surface area as body temperature increases, and temperature-dependent variation will be constrained at the upper end of the range of *cMR*, as predicted by the heat dissipation limit (HDL) theory of Speakman and Król (2010)^1^. These authors proposed that species encounter two kinds of ecological constraint on metabolic rate: when the food supply is limiting, competition will favor animals that have low *rMRs*, but when fuel energy is abundant, then a physiological constraint, specifically the maximal rate of heat dissipation, will limit the evolution of faster cellular *MRs*^1^.

## RESULTS

### Allometric scaling of vertebrate MRs by maximum likelihood

In conventional studies of the evolution of metabolic rate, constraints such as mass, temperature, aridity, and diet have been inferred from the slope and residual analysis of the regression of log *rMR* on log *M* (see, for example^14, 20, 69, 74, 75^). We first established that our sample of 20 vertebrate species in MirGeneDB was broadly representative of vertebrates with regard to the distributions of mass and *rMR*. As shown in Figure 1A, the *rMR*s of the 14 endotherms and 6 ectotherms exhibited a conserved relationship to mass, with allometric slopes (0.75 and 0.88 respectively) very similar to those reported in previous, large-scale studies^76, 77^. When *M*^exp^ was fit by maximum likelihood to the distribution of *rMR*^78^ (with resolution *M*^0.1^), the optimal regression yielded a common slope of 0.80 for both ectotherms and endotherms, with normally distributed residuals and a smaller residual error than the log-based method for either endotherms or ectotherms (*P* = 6.36e-32) (Figure 1B). Importantly, even though the variation of *rMR* with respect to *M*^exp^ is heteroscedastic (Figure 1B)^78^, when the scaling of *M*^exp^ is optimized in order to match the ratio of *rMR*/*M*^exp^ to the distribution of a third covariate, such as *miRNA.Fam*, the variation is homoscedastic (Figure 4 and Supplementary Figure S2), and therefore suitable for comparing the dependence of *cMR* on covariates across a wide range of species sizes^70^.

**Figure 1.**
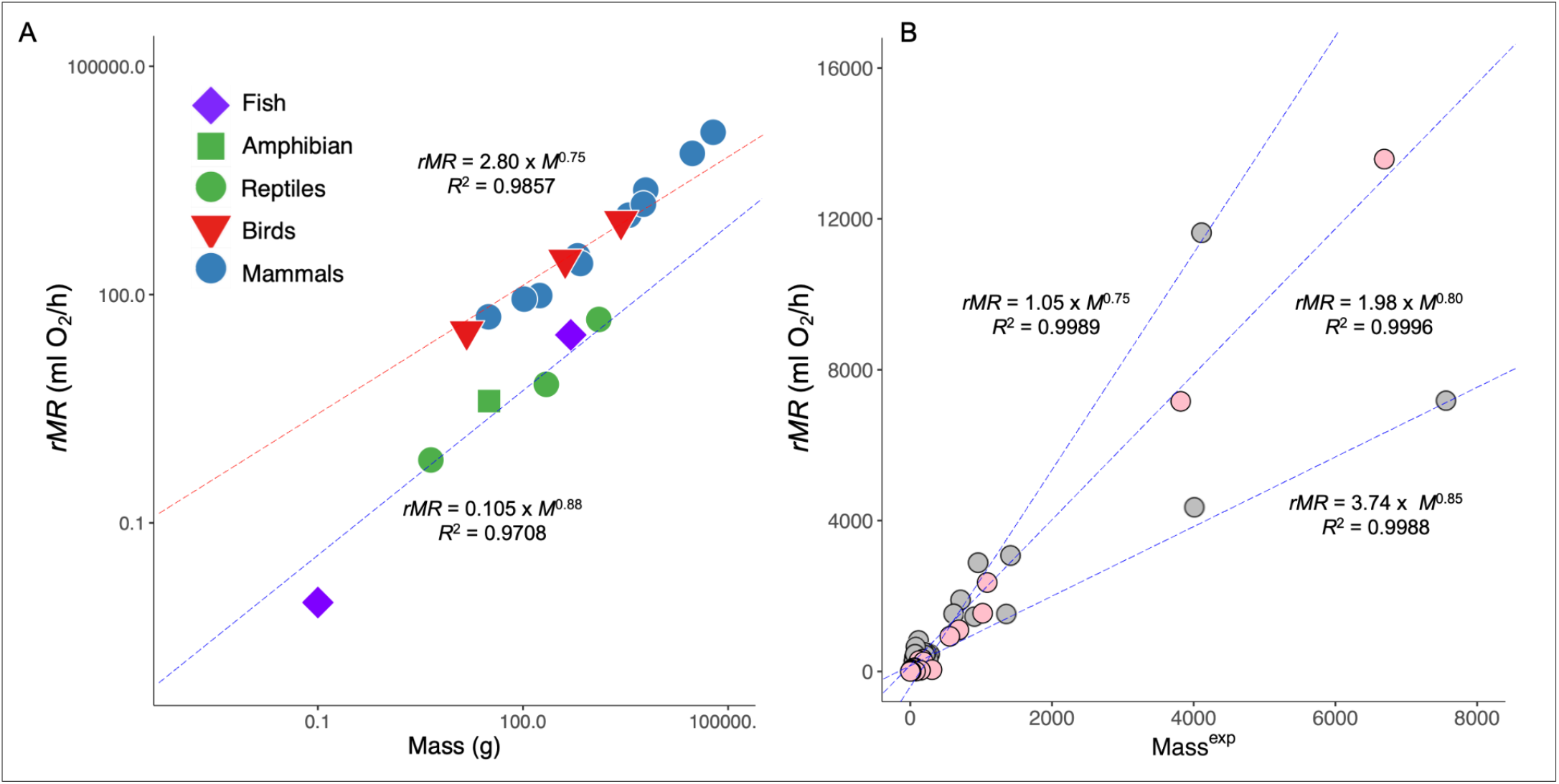
Comparison of the allometric scaling of the *rMR*s of 20 vertebrates by (A) the conventional power law method, or (B) maximum likelihood scaling according to the optimal fit (*M*^0.80^: pink circles) and two suboptimal fits (*M*^0.75^, *M*^0.85^: gray circles).

As shown in Figure 2, maximum likelihood scaling of *cMR* across the arithmetic range of *rMR* is more robust to variation in mass and more informative than the log-based method regarding the dependence of *cMR* on the ratio of surface area/mass, thus providing greater insight into the biophysical nature of the relationship. For example, when scaled by maximum likelihood, the optimal slopes of *cMR* with respect to *miRNA.Fam* for five early-branching mammals (mean *T*_b_ = 34.8°C) and five late-branching mammals (mean *T*_b_ = 37.6°C) were *M*^0.78^ and *M*^0.70^, respectively (*P* = 1.54e-02, ANCOVA) (Supplementary Table S3 and Supplementary Figure S3). However, when the corresponding *sMR*s were inferred from the slopes of log *rMR versus* log *mass* (*M*^0.75^ and *M*^0.74^, respectively) they were not different (*P* = 0.2240, ANCOVA). The greater resolution of the maximum likelihood approach can be attributed to the fact that the relationship between log *rMR* and log mass is governed by many interacting factors, while a single mechanism is sufficient to account for the dependence of *cMR* on *miRNA.Fam*.

**Figure 2.**
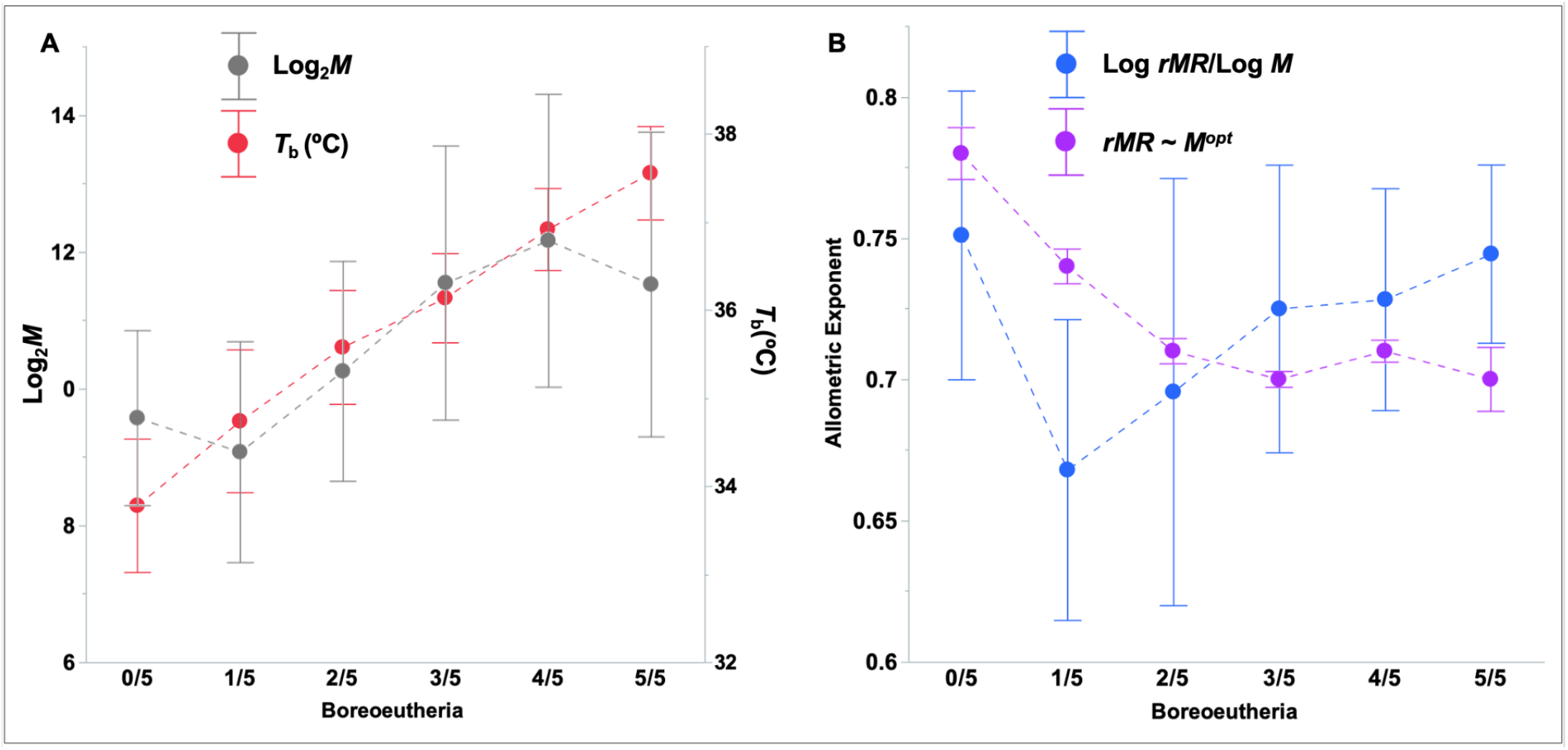
“Sliding Window” comparison of the inference of *sMR* and *cMR*. Staggered groups of five mammals ranked by *T*_b_ were designated by the number of Boreoeutheria. Panel A. The effect of group composition on the mean value of body temperature or mass (right axis). Panel B. The effect of group composition on the allometric exponent of *sMR* (the slope of log *rMR versus* log *M*) or the maximum likelihood exponent of *cMR* (means ± ½ standard deviation).

There was a significant anti-correlation between optimal *cMR* exponent and mean body temperature in samples of five mammals (*P* = 0.020): as mean *T*_b_ increased, the relationship between *miRNA.Fam* and *cMR* became more highly dependent on surface area. Furthermore, as predicted by the HDL theory, the optimal *cMR* exponent asymptotically approached a limiting value of 0.70. In contrast, the corresponding slopes of log *rMR versus* log *M* were highly variable and did not exhibit any consistent change with mean body temperature (Figure 2B).

### The acquisition of microRNA families accelerated in late-branching mammals

The number of families of conserved, microRNA-encoding genes (*miRNA.Fam*) increased steadily in the order of divergence of vertebrate clades, except among Archosauria (Figure 3A). The variation in *miRNA.Fam* was unrelated to genome size or the number of coding genes (Supplementary Table S3), but was 85% correlated with *cMR.75* (*rMR*/*M*^0.75^) across 16 non-avian vertebrates (*P* = 1.02e-07, Kendall’s rank correlation) (Figure 3B). The acquisition of shared microRNA families accelerated in placental mammals, with increases of 43 in the 64 million years between the divergences of Theria and Placentalia, and 27 in the 8.7 million years between the divergence of Atlantogenata and Euarchontoglires^79^, corresponding to rates of gain of 0.66% per million years in pre-placental theria *versus* 2.27% per million years in placentals. Consequently, the number of microRNA families common to the four Euarchontoglires in our dataset (152) is nearly twice the number shared by the Theria (82), as is the number of genes (Supplementary Table S1). Of these 152 conserved families, 13 have been shown to play a role in placental development or function^81^ but, at least in humans, no microRNA gene or family appears to be uniquely expressed in the placenta^67^.

**Figure 3.**
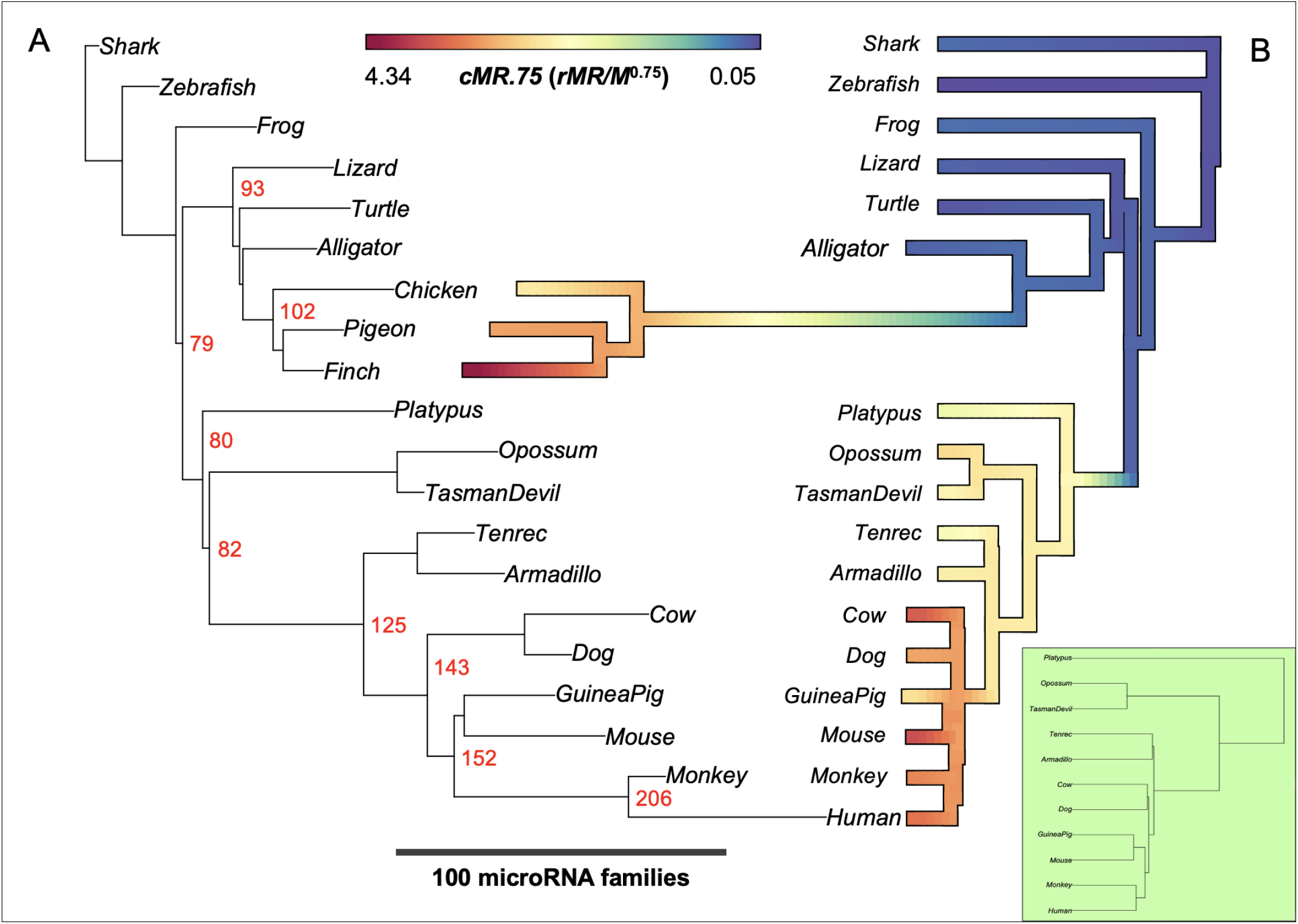
Evolution of the number of microRNA families and cellular metabolic rate (*cMR*). A. Distance tree of microRNA families. Branch lengths represent the total number of families. Node values represent the number of shared microRNA families in each clade. B. An ultrametric reference phylogeny (inset) was assembled based on divergence times drawn from Álvarez-Carretero et al 2022)^79^ for mammals and Irisarri et al (2017)^80^ for earlier-branching vertebrates. The reference tree was rescaled (BayesTraits) according to the rate of evolution of *cMR.75*, which corresponds to the optimal allometric scaling in a model of stabilizing selection of *rMR* with respect to the number of microRNA families in tetrapods and mammals (see Figure 5). Values of *cMR.75* inferred at ancestral nodes (BayesTraits) were then mapped onto this variable rate tree (see Methods).

The rate of evolution of *cMR.75* increased markedly at each stage in the bird lineage, in apparently inverse fashion to the number of microRNA families (Figure 3A). Among mammals, comparison of branch lengths between the variable rate tree for *cMR.75* and the reference constant rate tree (Figure 3: inset) indicated compression of the terminal branches and branches leading to the ancestral therian and placental mammal, while the branch leading to the ancestral Boreoeutherian was expanded, marked by a sharp increase in *cMR.75* (Figure 3B; for additional analysis, see Figure 7).

### The association between cMR and miRNA.Fam is independent of phylogenetic distance

Phylogenetic linear regression indicated that the distribution of *miRNA.Fam* across 14 non-avian tetrapods accounted for 84% of the variation in optimally scaled *cMR* (*rMR*/*M*^0.68^, *P* = 2.26e-06), independent of phylogenetic distance (Pagel’s *lambda* = 0.00) (Figure 4A and Supplementary Table S3). The optimal allometric slope, 0.68, was significantly different from 0.78, the slope of log *rMR versus* log *M* (*P* = 8.02e-03, ANCOVA). Similar results were obtained when only mammals were considered, which yielded optimal allometric exponents of 0.71 for *cMR* and 0.77 for log *rMR versus* log *M* (Supplementary Table S3). The variation of *cMR* with respect to the number of microRNA genes (*miRNA.Gen*) was weaker and, in the case of mammals, the association appeared to be secondary to phylogenetic distance (Pagel’s *lambda* = 0.617, Supplementary Table S3).

**Figure 4.**
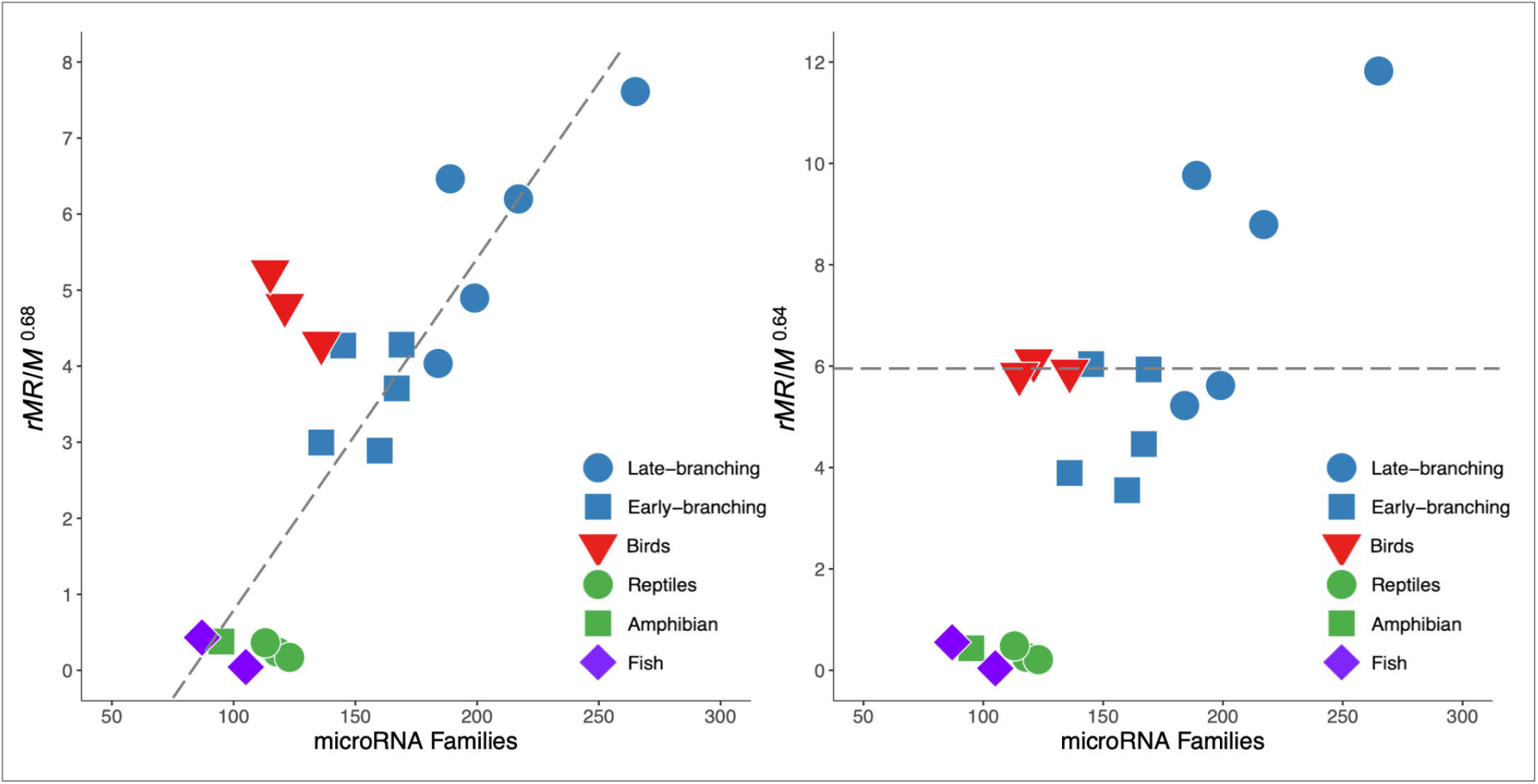
Phylogenetic regression of *cMR.68* (*rMR*/*M*^0.68^) on *miRNA.Fam*. A. The number of microRNA families in non-avian tetrapods accounted for 84% of the variation in *cMR*68, independent of phylogenetic distance (see Supplementary Table S2). B. The same data scaled to the slope of conventional *sMR* (log *rMR*/log mass = 0.64) in 211 birds^29^.

The previous analysis excluded the three bird species in our sample (chicken, dove and finch) since the absolute number of avian microRNA families decreases in phylogenetic order (Figure 3A). However in large datasets, the log-based allometric exponent for *rMR* in birds is markedly lower than that found in mammals, 0.64 *versus* 0.73^29, 69, 74, 76^. When scaled to *M*^0.64^, the slope of the regression of the bird *cMR* on the number of microRNA families was 0.00 (Figure 4B), and turned positive for any allometric exponent < 0.64. That is, when scaled with an allometric exponent < 0.64, bird *cMR*s did vary positively with *miRNA.Fam*. Thus, as in the case of mammals, the optimal allometric scaling of *cMR* with respect to *miRNA.Fam* in birds may be displaced toward a greater dependence on surface area *i.e.* lower allometric exponent, than is indicated by the slope of log *rMR versus* log *M* (Supplementary Table S3).

### Cellular MR variation in relation to the number of microRNA families conforms to an OU model of stabilizing selection

We compared two models for the evolution of cellular metabolic rates in relation to the microRNA complement: a Brownian motion (BM) process, where *cMRs* diverge as a function of phylogenetic distance, *versus* an Ornstein-Uhlenbeck (OU) process, according to which *cMR*s, under the influence of stabilizing selection, converge asymptotically toward the distribution of an independently varying predictor^8^, in this case, the number of microRNA families. We used two complementary approaches: first, we compared how well each model fit the relationship of *cMR* to *miRNA.Fam* across the range of allometric exponents 0.66 - 0.80; second, using the same phylogeny and distribution of microRNA families, we compared the actual distribution of *cMR* values with those generated by BM and OU simulations that included adaptive shifts at different stages of mammalian evolution.

For any allometric scaling of *cMR* in the range *M*^0.66^ - *M*^0.80^, the BM model for the evolution of *cMR* was inferior to the OU model (Figure 5A): the maximal ΔAICc (BM-OU) difference, found at at *M*^0.75^, was +2.68 before correction (Supplementary Figure S4) and +6.88 after empirical correction of the bias recorded in 40 OU and 40 BM simulations on the same phylogenetic tree (Supplementary Table S4). The inferred rate of adaptation of *cMR* with respect to *miFam* at *M*^0.75^ corresponded to 12.63% of the tetrapod tree (Figure 5A), or 4.92% when only the 10-mammal tree was considered (Figure 5B). Cellular *MR*s exhibited a similar ΔAICc maximum with respect to the number of microRNA genes (*miRNA.Gen*), however no such inflection was observed in models of *cMR* variation in relation to genome size (*Gnm*) or number of coding genes (*Cdg*), where AICc values declined monotonically with increasing allometric exponent (Supplementary Figure S4).

**Figure 5.**
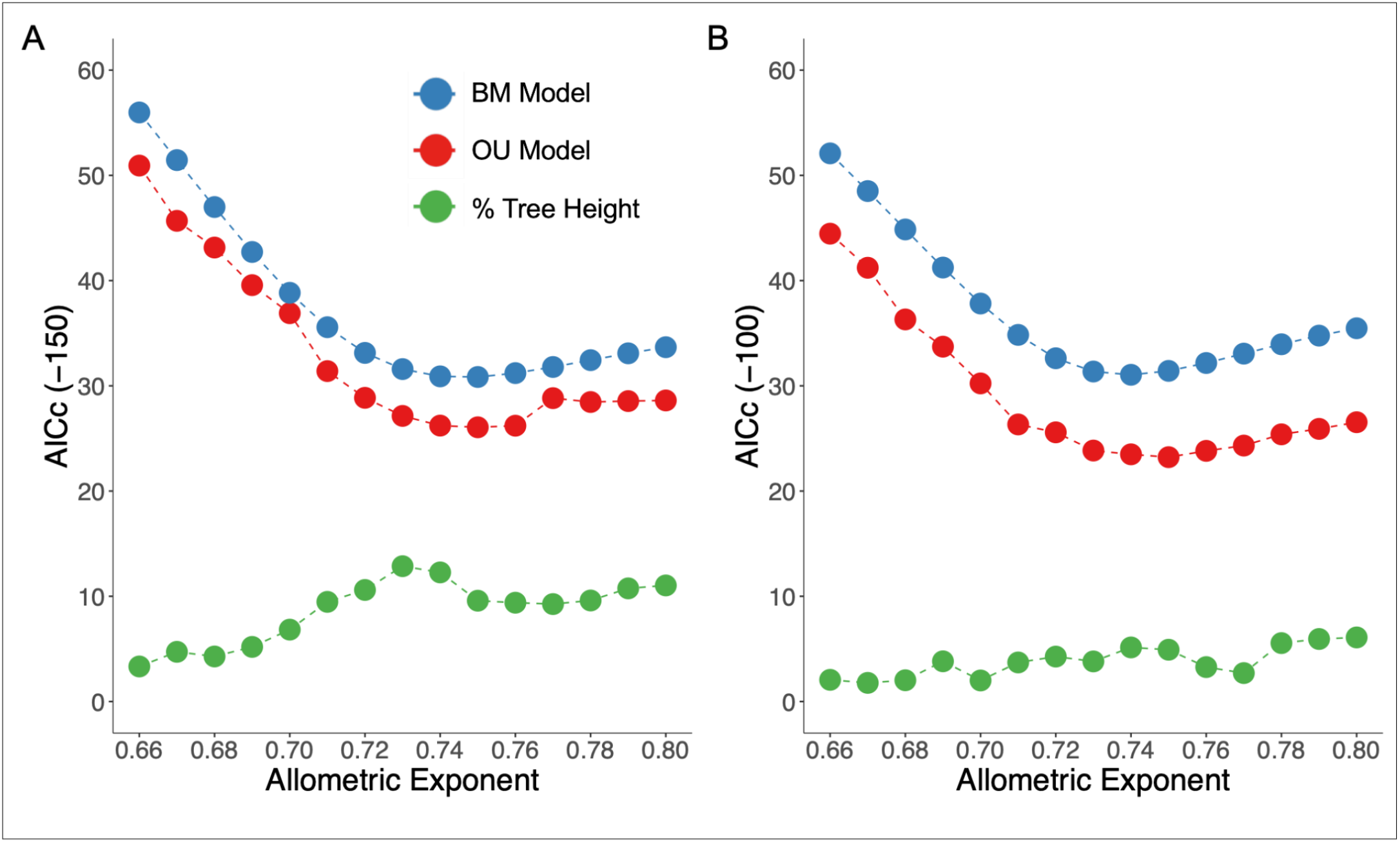
BM and OU models for the variation of c*MR* with respect to *miRNA.Fam.* Curves represent the corrected values of the Akaike information criterion (AICc) for each model, and the rate of adaptation (% tree height) for the OU models (see Methods). Note the difference in ranges of the ordinate axes. A: 14 tetrapods (excluding birds and cows). B: 10 mammals (excluding cows).

In our dataset there was an abrupt increase in the number of microRNA families between the two Atlantogenata (Tenrec: 160, Armadillo: 169) and the two Laurasiatheria (Dog: 184; Cow: 212), suggesting the emergence of a new optimum in the adaptive landscape^2, 3^, ^4^. We therefore simulated the evolution of *cMR* with respect to *miRNA.Fam* in 14 tetrapods by either a BM or OU process, incorporating two adaptive shifts^10^, the first at the divergence of mammals, and the second shift corresponding to the divergence of either Placentalia, Boreoeutheria, or Euarchontoglires (Figure 6A). The best model, an OU process with an adaptive shift at the divergence of Boreoeutheria, accounted for 95% of the variation in *cMR.75* (*P* = 4.10e-09), compared to 88% for the corresponding BM model (Figure 6B). Estimates of the rate of adaptation increased in the order of the phylogenetic position of the second adaptive shift, in other words, decreasing distance from the tips of the phylogenetic tree (*P* = 5.19e-06, ordinal logistic regression) (Figure 6C). In sum, for both the OU and BM models of the variation in *cMR* across mammals, the optimal allometric scaling with respect to *miRNA.Fam* matched the conserved ¾ power law model for the overall dependence of *rMR* on mass in mammals. It remains to be determined whether in birds, by analogy, the optimal allometric scaling in BM and OU models of *cMR* dependence on *miRNA.Fam* corresponds to the ∼ ⅔ power law model that fits avian resting metabolic rates^29, 69, 74, 76^.

**Figure 6.**
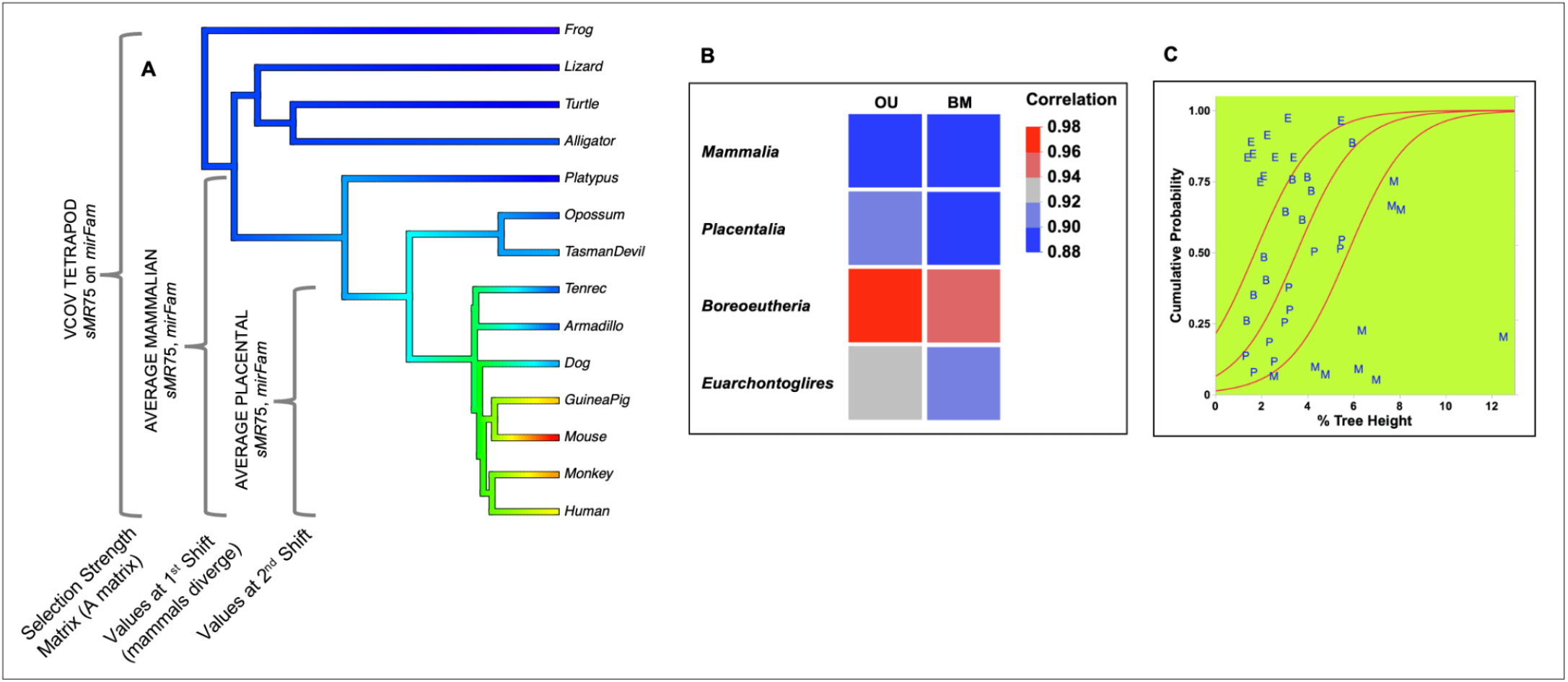
Design and analysis of simulated adaptive shifts in the evolution of *cMR* in relation to the predictor *miRNA.Fam*. A. The variance/covariance matrix for the regression of *cMR.75* on *miRNA.Fam* was used as the selection strength matrix to generate pairs of values for these traits. The average values of these traits across all mammals, or across placental mammals, were assigned to the first shift (divergence of mammals) and second shift, respectively. Phenogram colors correspond to the log of the *cMR.75* values scaled to the original, 36-fold range. B. Correlations with the actual *cMR.75* distribution of the average of ten simulations of the OU or BM process in 14 tetrapods. Adaptive shifts were modeled at the divergence of Mammalia (single shift), with a second shift at the divergence of either Placentalia, Boreoeutheria or Euarchontoglires (*P* = 1.23e-05 for the difference in correlation between OU and BM for 10 simulations with the Boreoeutheria shift). C. Validation of the estimated rates of adaptation: prediction of shift positions according to the % tree height obtained from ten OU models. Shift positions are indicated by the first letter of each diverging clade (M, P, B, E; *P* = 5.19e-06, ordinal logistic regression).

### The divergence of Boreoeutheria was associated with a step increase in the rates of evolution of *cMR.75, miRNA.Fam and body temperature*

Based on the evidence that the dependence of *cMR* on *miRNA.Fam* across mammals is sensitive to temperature and underwent an adaptive shift, we predicted increases in the branch-wise rates of evolution of all three traits. Using a Bayesian variable rates algorithm, we generated phylogenetic trees in which branch lengths were scaled in proportion to the rate of evolution of each trait, relative to an expectation of a Brownian motion process<u>^7,^</u> ^73^ (Figure 7).

**Figure 7.**
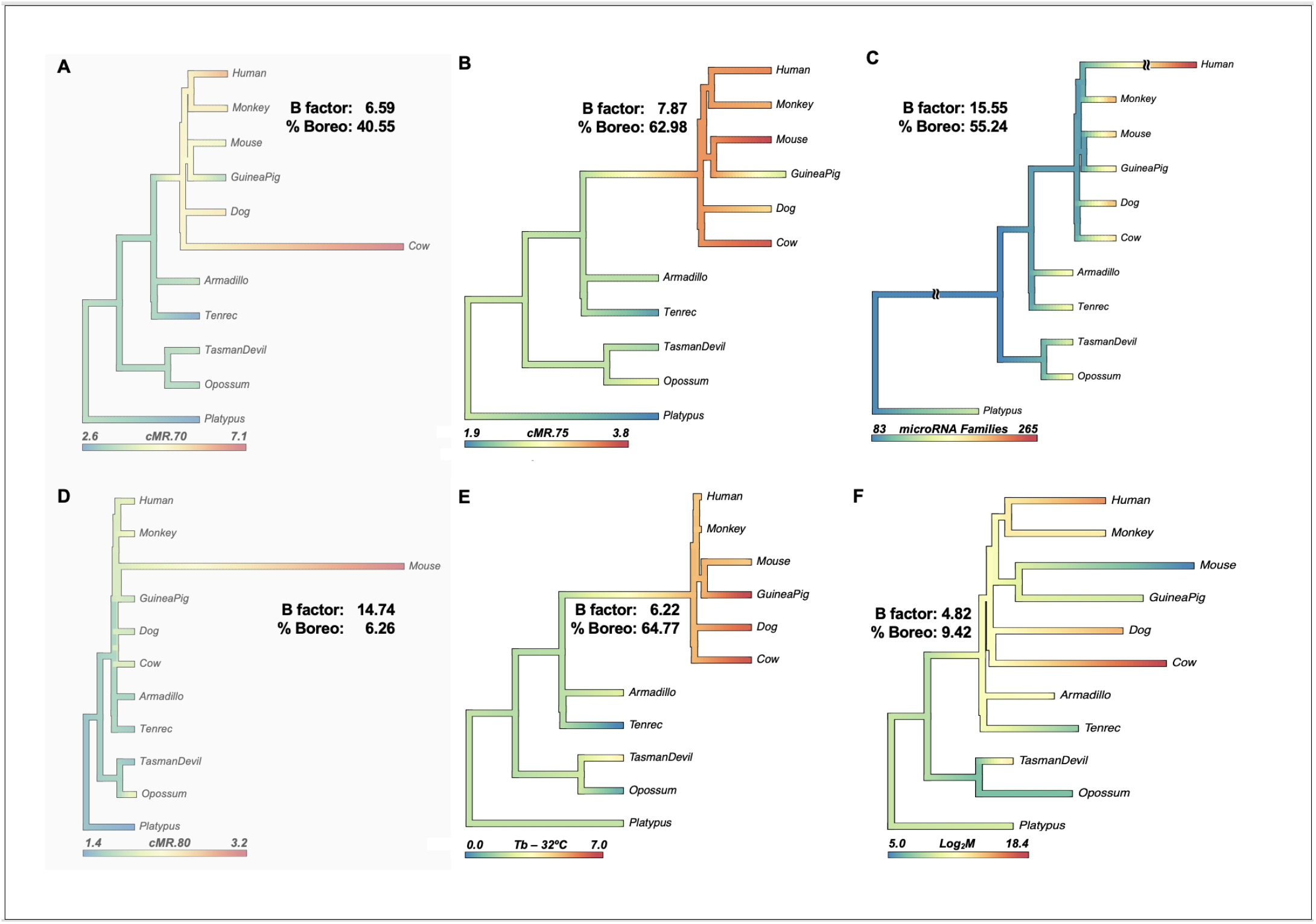
The ultrametric reference phylogeny of mammals (constant rate model) was rescaled according to the rate of evolution of each of six traits (A - F): *cMR.70*, *cMR.75*, *cMR.80*, *miRNA.Fam*, *T*_b_(°C) and log_2_*M*. The length of the branch for the divergence of Boreoeutheria was compared (%) to the distance between the divergence of Theria and the divergence of Primates. Bayes factors were computed by comparing the median log likelihoods of the 100 most likely trees to the median log likelihoods of the trees with the smallest non-zero sum of changes to branch *i.e.* those trees that differed the least from the constant rate model. This criterion was adopted in lieu of the marginal likelihoods of the variable and constant rate models, which reflect the overall fit of all of the branches. The range of ancestral node values inferred in a Brownian motion context were mapped onto each tree (see Methods). The value at each node represents the median of the ten reconstructions with the highest likelihood. In panel C (*miRNA.Fam*), the lengths of the branches leading to humans and to the ancestral Therian have been reduced by half.

As shown in Figure 7 panels A, B and D, arbitrarily changing the allometric exponent changes the relative magnitudes of the branch lengths of the variable rate trees for *cMR*: lower slopes (lower exponents/higher ratios of *rMR*/*M*^exp^) magnify the difference at the upper end of the range of mass, and *vice versa*. Based on comparison of the median log likelihood of the 100 trees with the greatest (1%) likelihood and the 100 trees with the smallest (1%) sum of adjustments to branch lengths, *i.e.* those trees most similar to the reference constant rate model, the variable rate trees representing *cMR.75*, *miRNA.Fam* and *T*_b_ (Bayes factors > 6) and log_2_ mass (Bayes factor > 4) were strongly supported^73, 82^. In the variable rate trees for *cMR.75*, *miRNA.Fam* and *T*_b_, the length of the branch representing the relative rate of evolution between Atlantogenata and Boreoeutheria was 6 - 7 times the length of the corresponding branch in the tree representing the evolution of log_2_ mass (Figure 7). Moreover, for these three traits, the branch lengths representing the descent of Primates from Theria were 99% correlated (Supplementary Table S5). If the variable rate tree for *sMR.75* depicts the outcome of selection, then the remaining phylogenetic variance ought to reflect only genetic drift, with minimal difference in credibility between the adaptive OU and random BM models, as was the case: ΔAICc (BM-OU) at *cMR.75* decreased to −0.53 (before correction) (Supplementary Figure S5).

Losses have been rare in the evolution of microRNA repertoires^83^, and we noted only gains in the terminal branches of the *miRNA.Fam* tree, with the human lineage exhibiting an exceptional increase (Figure 7C). On the other hand, accelerated rates of size evolution and marked decreases in body size were inferred in the four smallest mammals: opossum, tenrec, mouse and, to lesser extent, guinea pig (Figure 7F). The adaptations that accompanied these reductions in size differed between the two permanent homeotherms (mouse, guinea pig) and the two heterotherms (opossum, tenrec). In the homeotherms, the adaptations suggested thermoregulation to limit maximal heat generation: in the mouse, a severe reduction in size and increase in the surface area/volume ratio appears to have facilitated an increase in cellular *MR*, but not *T*_b_. In the guinea pig, with 20 times the mass of the mouse, an increase in body temperature appears to have been offset by a reduction in cellular *MR*. (We have assumed that mouse and guinea pig have similar thermal conductances.) In contrast, the two heterotherms exhibited concurrent reductions in both body size and temperature, indicative of a limited energy supply, but opposite changes in *cMR* (Figure 7E, B). The interpretation of these responses is complicated by the fact that both opossums and tenrecs exhibit daily torpor^15, 18, 84^.

Since the acquisition of microRNAs and increases in *cMR* in mammals were generally associated with higher body temperatures and larger body sizes (Supplementary Table S1)^85^, we used two approaches to further delineate the interdependence among these traits. Conventional analysis of covariance indicated that the dependence of *cMR.75* on *miRNA.Fam* in mammals was still significant after controlling for mass (*P* = 0.0267), but not after controlling for body temperature (*P* = 0.2133) (Supplementary Table S6). Bayesian regression confirmed that the inclusion of *T*_b_ improved the overall validity of the model, while in the model including mass, there was a lack of support for the regression coefficient for (log_2_) mass (Supplementary Table S6).

### Faster sMRs and homeothermy in late-branching placental mammals are associated with the number, not kind, of predicted microRNA-target interactions

Of the 146 microRNA families common to all six Boreoeutheria in this study, 19 were not shared by both Atlantogenata (tenrec and armadillo): 10 were present only in tenrec, 8 were present only in armadillo, and only a single microRNA family, miR-370, was absent from both genomes (Supplementary Table S7). Moreover, for three of these 19 microRNAs there were no conserved target sites: miR-105, miR-370 and miR-6715. It cannot be excluded that the difference in energy regimes between early- and late-branching placental orders is related to the activities of some combination of the remaining 16 microRNAs, affecting targets unique to, or differentially expressed in later-branching placental mammals. Any such discovery, however, would not shed light on the central question of this study, the nature of the broad correlations across mammals between cellular metabolic rate, the size of the microRNA repertoire, and the predicted number of microRNA-target interactions.

We assume that global repression of translation by the combinatorial activity of microRNAs is proportional to the number of microRNA-target interactions and target sensitivity to repression. Since the TargetScan database lacks data from Atlantogenata, we tested this assumption by comparing the predicted level of global repression across frog, chicken and four late-branching placental mammals (all Boreoeutheria), relative to that of a marsupial (opossum). Based on the number of conserved 3’-UTR sites (TargetScan database^63, 86^) of genes targeted by conserved microRNAs (MirGeneDB^67^), *i.e.* those sites most likely to have been retained by natural selection, we evaluated the following model to predict the energy cost saved by the sum of conserved microRNA-target interactions per cell:

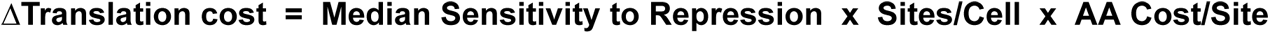

Given that the cost of protein synthesis accounts for ∼20% of cellular metabolic rate, we predicted that the ∼ 30% increase in *cMR.*75 between marsupials and late-branching placental mammals would be associated with an increase in overall microRNA-mediated repression of translation of ∼ 150%. Figure 8 depicts the ratios of the value for each factor in the model to the corresponding value in the opossum. Relative to the opossum, the proportion of coding genes targeted and the number of 3’-UTR sites/mRNA were both approximately 50% higher in the placental mammals (Supplementary Tables 2). The total number of predicted microRNA-target site interactions per cell increased 100%, 140% and 250% in dog, mouse and human while the corresponding median values for sensitivity to repression by combinatorial microRNA activity (*C*_n_) decreased by 11%, 12% and 16.7% (Figure 8). After taking into account the species distributions of target proteins 500 - 5,000 AA in length, the predicted ratios of overall translation cost were 180% (dog), 192% (mouse) and 269% (human) (Figure 8). These results are in reasonable agreement with our prediction, considering that we made no correction for the relative abundance of each protein, and that the *C*_n_ values have likely been overestimated, as they are based on the most effective microRNA from each family, ^63, 86^.

**Figure 8.**
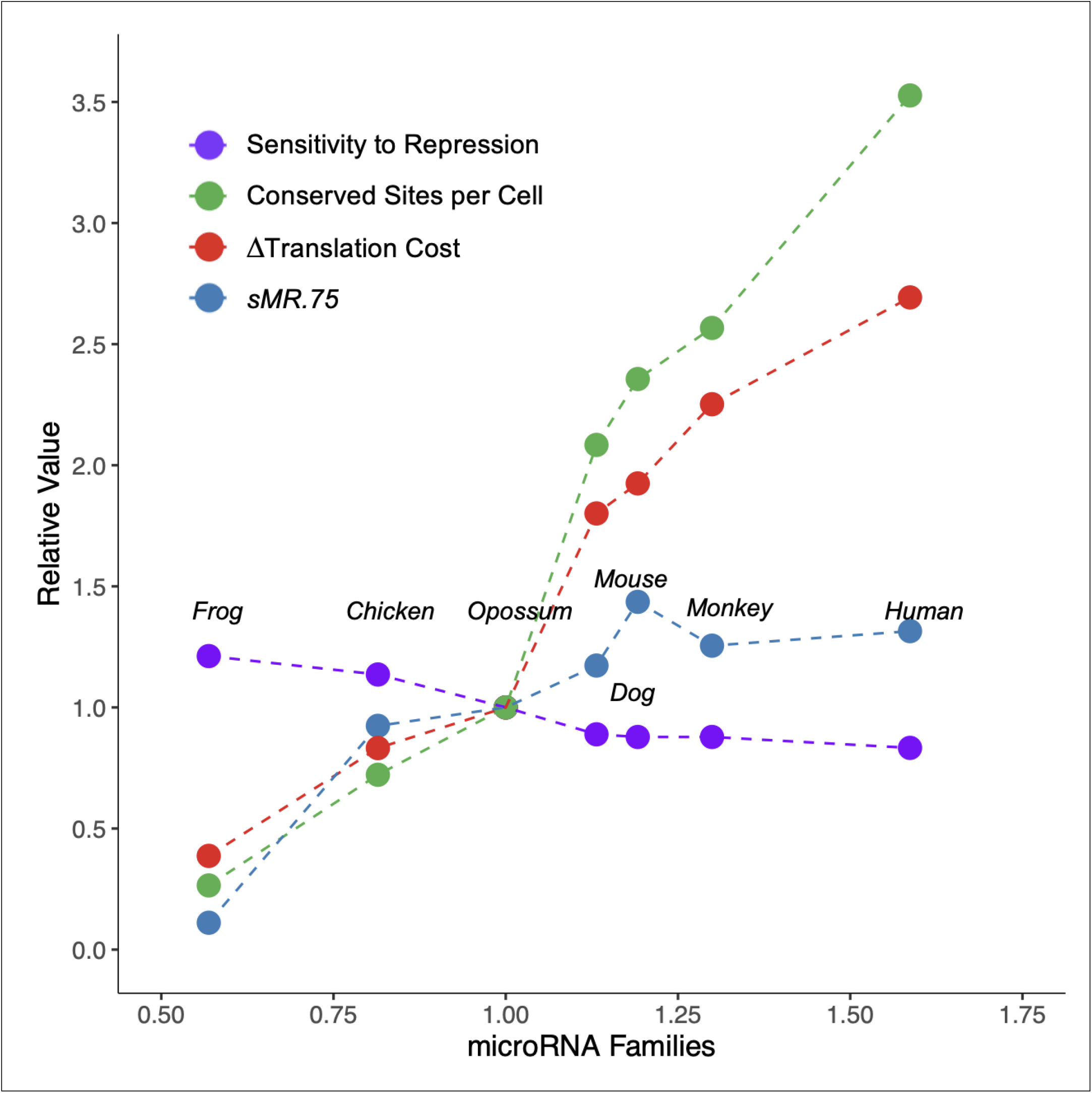
Evolution of factors in a model for energy cost saving by global microRNA activity. *Sensitivity to Repression*: the median value of the distribution of cumulative weighted context scores for all models of the conserved targets of conserved microRNAs (see Methods). *Conserved Sites per Cell*: the total number of sites was standardized with respect to the number of coding genes in each species. *AA cost/site*: the weighted sum of 500 amino acid-length bins for the distribution of a non-redundant set of target proteins between 500 - 5,000 amino acids (see Methods).

## DISCUSSION

### Variable allometric scaling: a new method to explore the evolution of cellular MR

Given the high metabolic cost of protein synthesis and its strict relationship to cell size^32, 35^, we assumed that global attenuation of intrinsic translation noise is energy-saving and proportional to both cellular *MR* and the size of the microRNA repertoire. Using variable allometric scaling in conjunction with phylogenetic comparative methods, we estimated *cMR* by fitting the *rMR*/*M*^exp^ ratio to the number of microRNA families. This report thus complements recent studies that have inferred cellular *MR* from separate analyses of the variation of log mass and log *rMR*^27, 72, 85^. The greater resolution of the arithmetic range of *rMR* for fitting *rMR*/*M*^exp^ to *miRNA.Fam*, together with this method’s robustness to variation in mass, made it possible to detect changes with body temperature in the surface area dependence of the *cMR*-*miRNA.Fam* relationship, even among small groups of mammals.

### MirGeneDB is a new reference for comparative biology research

Previous comparative studies have referenced molecular traits, such as genome size, the number of protein-coding genes, or the quantity of non-coding DNA (for example, Choi et al, 2020^87^). With the availability of the manually curated microRNA gene database, MirGeneDB, we are able to extend this approach to conserved microRNA families and genes in a wide range of vertebrate species and organs^67^. A major concern in microRNA research in the past has been the quality of the online repository miRBase. This database contains many false-positives while at the same time missing a large number of *bona fide* microRNAs from undersampled species, which were misinterpreted as secondary losses^88, 89, 90, 91, 92, 93, 94, 95, 96, 97^. Consequently, the conservation and utility of microRNAs as phylogenetic markers has been questioned in the past^98^. However, these omissions were clearly technical artifacts, since losses of microRNA families have rarely been observed in well-annotated repertoires^67^. We and others have recently successfully employed MirGeneDB complements in a range of comparative^99^, phylogenetic^65, 100, 101^, developmental^80, 102, 103^ and evolutionary studies^104^, as a validation cohort for experimental findings^105, 106^, and as a reference for the development of MirMachine, a highly accurate machine learning-based tool for the automatic annotation of microRNA families^107^.

Exhaustive sequencing and genome curation have revealed that the number of families of conserved, microRNA-encoding genes has grown steadily over the course of vertebrate evolution, with very low rates of loss^67, 83^. Notably, the acquisition of shared microRNA families and conserved 3’-UTR target sites accelerated in placental mammals, doubling the potential number of conserved microRNA-target interactions (Supplementary Tables S2). The acquisition of microRNA families and targets coincided closely with an elevation in *rMR*, body temperature and propensity for homeothermy among those mammalian orders that diverged more recently than Chiroptera^19, 20^: out of six mammalian orders with mean body temperatures significantly higher than that of bats, the only clade that diverged earlier than Chiroptera, Macroscelidea (Supplementary Figure S1), was a lineage characterized by an extreme reduction in size^12^.

Conversely, out of eight orders with lower average body temperatures, the only one that diverged later than Chiroptera, Pholidota, is characterized by species with relatively large masses, low *MR*s and body temperatures close to ambient^108^ (Supplementary Figure S1), suggesting a limited capacity to dissipate heat^1^. Since the order Pholidota has undergone relatively few speciation events since the Cretaceous^109^, this observation may be a reminder that the evolution of faster rates of metabolic heat generation depended on opportunities for the evolution of faster mechanisms of heat dissipation^28, 29^.

### Cellular MR variation with respect to miRNA.Fam fits a model of stabilizing selection

For a BM process, between-species trait covariances are proportional to their shared ancestry, and a BM model will be favored if most of the crown species diverged recently relative to the total age represented by the phylogeny, as is the case of the mammalian tree used in this study^78^. Furthermore, for a small sample and phylogeny the mvSLOUCH algorithm for the comparison of OU and BM models exhibits a bias towards BM that has been attributed to a difference in the effective population size of each process^110^. For example, in a re-analysis of Darwin’s data for 13 species of finch^111^, measures of effective population size yielded correction factors of ∼ 0.5 and ∼ 0.9 for the BM and OU models, respectively^110^. However, even before applying an empirical correction for bias, we found that an OU process was consistently favored over a BM process as a model for the evolution of *cMR* with respect to *miRNA.Fam*, in both our 14-tetrapod and 10-mammal datasets.

Assuming constant rates of evolution, the optimal slopes obtained by phylogenetic linear regression for the dependence of *cMR* on *miRNA.Fam* across tetrapods or mammals were *M*^0.68^ and *M*^0.71^, respectively (Supplementary Table S3). However, the AICc information criteria for both BM and OU models reached their minimal values when *cMR* was scaled to *M*^0.75^, where BM and OU processes most precisely simulated the biphasic evolution of *cMR* (Supplementary Figure S6). This example illustrates an aspect of the scaling of *MR* that is generally overlooked when comparing allometric exponents inferred from the slopes of log *rMR versus* log *M*, namely that a differences in allometric slopes generally represents a shift in the rank order of the ratios between *sMR* (or *cMR*) and a third covariate. For example, among the 10 mammals included in this study, the dog has the highest ratio of *cMR* to *miRNA.Fam* when *cMR* is scaled to *M*^0.70^, second highest at *M*^0.75^ and fourth lowest at *M*^0.80^.

BM and OU models of the variation in number of microRNA genes exhibited the same *M*^0.75^ optimum, albeit in a higher range of AICc values, but no optimum was observed for *cMR* variation in relation to genome size or the number of coding genes (Supplementary Figure S4). Thus, *cMR* variation was most immediately linked to the number of microRNA families. Given that the mechanism by which microRNAs conserve metabolic energy is well established, the specific association between *miRNA.Fam* and *cMR.75* supports our assumption that global microRNA activity is proportional to the number of microRNA families and evolved under similar constraints. According to this conjecture, the relative rates of translation of low- and high-abundance transcripts, which is dictated by global microRNA activity^51^, ought to be energy efficient, as has been demonstrated previously^50^.

Three-quarter power scaling corresponds approximately to the conserved slope of log *rMR versus* log mass reported for mammals in countless reports since Kleiber (1932)^112^ and Brody and Proctor (1932)^113^ (see also Savage et al, 2004^114^). The present study indicates that this allometry also represents the optimal ranking of the ratios of *cMR/miRNA.Fam* and implies that the conserved ∼ ¾ power allometry is the outcome of multivariate selection primarily on the efficiency of cellular metabolism, and only secondarily on mass or total *rMR*^72^. This comports with our observation that *cMR.75*, *miRNA.Fam* and *T*_b_ underwent concurrent 6- to 7-fold increases in evolutionary rate between the divergence of Atlantogenata and Boreoeutheria, independent of the rate of evolution of mass: an increase above the background rate of evolution of 2-fold or more is considered evidence of positive phenotypic selection^73^. An implication for future studies is that the allometric relationship between *rMR* and mass obtained by fitting *cMR* to *miRNA.Fam* may be more relevant to the evolution of endothermy than the power law model inferred from the slope of log *rMR versus* log *M*. Such a comparison may now be performed on any set of genomes using MirMachine, which annotates microRNA families with a mean accuracy of 92% in Deuterostomes^106^, relative to the manually-curated database MirGeneDB^67^. For example, we have been able to confirm and extend the recent report of a step increase in the number of microRNA families in Simiiformes^115^: the range of *miRNA.Fam* detected by MirMachine in Catarrhini (197 - 208, n = 12) did not overlap the ranges in Platyrrhini (172 - 177, n = 4) or Strepsirrhini (164 - 181, n = 17) (Supplementary Table S8).

### Optimally scaled MRs may also predict the microRNA complement in birds

In linear phylogenetic regressions, the optimal *cMR* scaling with respect to *miRNA.Fam* in tetrapods (*M*^0.68^), mammals (*M*^0.71^) and Boreoeutheria (*M*^0.70^) exhibited greater dependence on surface area than did the log-based dependence of *rMR* on mass (*M*^0.78^, *M*^0.77^ and *M*^0.74^, respectively), an observation that may shed light on the correlation between vertebrate morphological patterns and microRNA complement size^116^. These authors demonstrated that morphological types, a function of aspect ratios, were significantly related to the number of microRNA families, but not to genome size or diversity of protein domains. Furthermore, the morphospace occupied by birds was significantly displaced from the reptilian-mammalian axis^116^, an interesting parallel to the shift in *cMR*-*miRNA.Fam* relationship that we have predicted in birds, where *cMR*s are positively related to *miRNA.Fam* only when the allometric exponent is set lower than ∼0.64, corresponding to the avian slope of log *rMR versus* log *M*^29, 69, 73, 75^.

Another factor that will impact the allometric scaling of *cMR* is the relative metabolic cost of protein synthesis. Muramatsu (1990)^117^ compared specific rates of protein synthesis in the chicken and a range of domestic mammals, all homeothermic Boreoeutheria. After correcting for egg production (14% of total protein synthesis^118^, 3% of *bMR*^119^) the specific rates of protein synthesis in chicken were 40 - 78% higher than in mouse, rat or rabbit, while the specific *MRs* of chicken were only 17 - 44% higher (Supplementary Table S9). If *miRNA.Fam* in birds is calibrated to a greater fraction of cellular *MR* than in mammals, then maximum likelihood fitting of the two distributions should yield a lower optimal allometric exponent, hence a higher ratio of *rMR*/*M*^opt^.

There is an important difference between birds and mammals in the regulatory context in which microRNAs operate. Ancestral birds underwent a reduction in genome size, from ∼ 2 Gb to ∼ 1 Gb, primarily by jettisoning transposable elements (TEs)^120, 121^. The massive loss of TEs in the ancestral avian genome contrasts with the steady evolutionary accumulation of TEs in mammals, where, with the exception of bats, they occupy 40 - 50% of the genome^122^. Retrotransposon sequences have been identified in the 3’-UTRs of 25% of mammalian coding genes and serve as binding sites for inhibitory KRAB-ZnF transcription factors^123^. The loss of TEs in the ancestral bird genome was accompanied by the nearly complete loss of KRAB-ZnFs^124, 125^, implicating microRNA activity as the principal mechanism for the downregulation of gene expression. Interestingly, bat genomes are also characterized by smaller size, loss of transposable elements^121, 126^ and shorter introns^127^ relative to non-volant placental mammals, suggesting that these reductions represent adaptations for the faster rates of signaling and energy expenditure associated with flight^128, 129, 130, 131^. For example, shorter introns enable faster mRNA splicing^132^, while compact genomes and nuclei, if found in smaller cells with a higher ratio of surface area/mass, would permit more rapid exchange of nutrients, as well as a higher neuron density in the brain^133, 134^.

### The predicted energy saved by global microRNA activity increased faster than cellular metabolic rate in placental mammals

The classification of conserved microRNA genes in MirGeneDB is congruent with the classification of conserved 3’-UTR target sites in TargetScan database^135^, but the latter lists targets for microRNAs that are excluded from the reference database, MirGeneDB^67^. For example, of the 265 thousand sites listed for the human in the conserved site database, only 173 thousand correspond to the reference microRNAs listed in MirGeneDB (Supplementary Tables S2). In order to exclude false positives, we considered only the conserved targets of these reference conserved microRNA families (see Methods).

For each species *C*_n_, the cumulative combinatorial repression per site, was estimated from the median value of the distribution of “cumulative weighted context++ scores”^63^. The CWCS model in the current version of TargetScan (.v8) accounts for 37% of the variation in messenger RNA levels following microRNA transfection of Hela cells, and probably overestimates the repression since, for each site, only the interaction with the most effective member of a microRNA family is evaluated^86^. After controlling for the number of coding genes in each species and the distributions of the lengths of the targeted proteins, the estimated energy cost saved by global microRNA activity in placental mammals was double that found in the opossum, consistent with the ∼ ⅓ increase in *cMR.75*. This is only a first-order approximation, as we did not try to account for the relative abundance of each protein. This agreement suggests an immediate energy-saving benefit proportional to the overall number of microRNA-target interactions, notwithstanding the lack of a scorable phenotype, which comports with evidence that the majority of mammalian-specific target sites appear to be evolving neutrally^136^.

### An adaptive shift at the divergence of Boreoeutheria simulates the evolution of cMR

The advent of placentation allowed some orders to exploit high quality resources through an increase in energy expenditure and reproductive output, adaptations denied to marsupials with access to the same resources^14^. This divergence in energy regimes was associated with differences in the relative prevalence of homeothermy *versus* heterothermy or torpor. Heterothermy is achieved by lowering the *T_b_*set point and evolved as an energy-saving strategy in small (< 1 kg) birds and mammals, notably bats, that are reliant on foraging and vulnerable to changes in food supply. After accounting for mass, climate and geography, earlier-branching placental clades, including Afrotheria, Xenarthra and Chiroptera, generally exhibit *T*_b_ < 36°C, a higher incidence of heterothermy, a lower proportion of skeletal muscle mass and lower metabolic rates, whereas homeothermy and *T*_b_ > 36°C predominate in Laurasiatherian orders that diverged after Chiroptera (excepting Pholidota) and Euarchontoglires (excepting Eulipotyphla and some Strepsirrhini) (Supplementary Figure S1). ^16, 17, 18, 19, 20, 21, 137, 138, 139^.

Based on the present range of species in MirGeneDB^67^, our demonstration that the *cMR* distribution in mammals was best simulated by an OU process with an adaptive shift at the divergence of Boreoeutheria supports a model with distinct adaptive optima for low- and high-*T*_b_ mammals. Figure 9 illustrates the estimated temperature gradients and optimal allometric exponents for the *cMR*-*miRNA.Fam* relationship in five early-branching, low *T*_b_ mammals (including Atlantogenata) and five later-branching, high *T*_b_ mammals (Boreoeutheria) (Supplementary Table S3). A separate optimum is shown for birds, with a predicted allometric exponent shifted to below the allometric slope (0.64) of log *rMR versus* log mass^29, 73^. The temperature gradients maintained by the low- and high-*T*_b_ mammals differ by approximately 8°C^108^ (Supplementary Figure S7), corresponding to a theoretical difference of 2.5% in thermodynamic efficiency (“Carnot Efficiency”) and potentially offsetting some of the additional cost of faster cellular *MR*s (see Methods for data sources).

**Figure 9.**
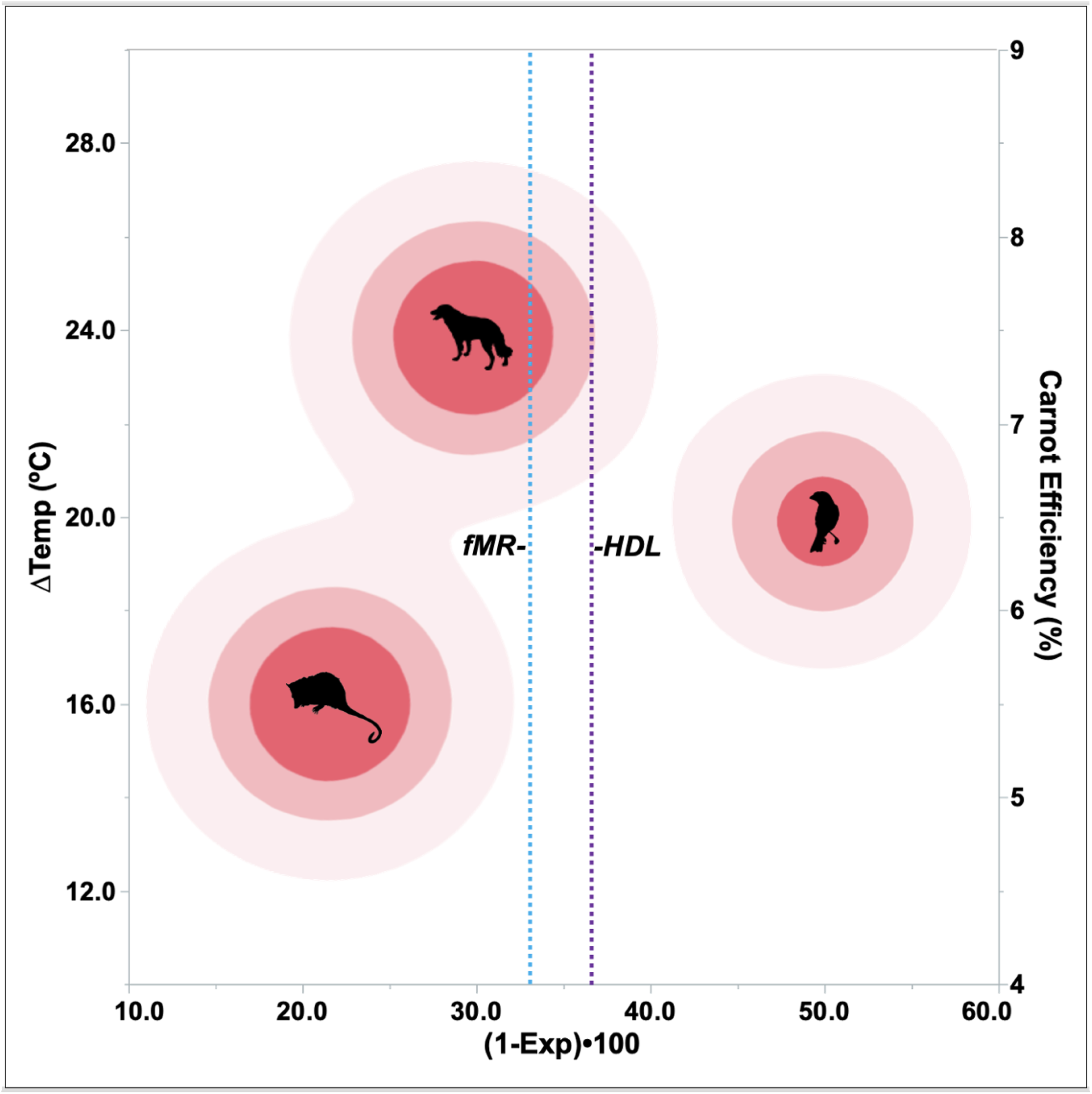
Contour plot of a hypothetical fitness landscape mapped on the relationship between the core-surface temperature gradient (**ΔTemp**) and **Exp**, the optimal allometric exponent for fitting *cMR* to the distribution of *miRNA.Fam*. The dashed lines represent the log-based allometric exponents for field metabolic rate (*fMR*) and the theoretical heat dissipation limit (*HDL*) in mammals (Speakman and Król, 2010)^1^. Silhouettes represent Boreoeutheria, earlier-branching mammals, and birds. (Phylophic © 2023: Marsupial, Sarah Werning; Dog, Tracy A. Heath; Bird, T. Michael Keesey). See Methods for data sources.

The inference of distinct adaptive peaks is based on the present range of mammalian microRNA repertoires in MirGeneDB, which represents only two of the four possible combinations of ancestry and phenotype: earlier-branching mammals with low *T*_b_’s and later-branching placentals with high *T*_b_’s. The addition of definitive microRNA repertoires to intermediate branches of the mammalian tree, such as those of low *T*_b_ Eulipotyphla, Chiroptera and heterothermic Strepsirrhini^21, 139, 140^, may eventually bridge these hypothetical optima and conform better to a biased random walk model that accommodates an abrupt change in evolvability^12^ related to the size of the microRNA repertoire. For example, among those orders of Laurasiatheria and Euarchontoglires where heterothermy is prevalent, *i.e.* Chiroptera and Strepsirrhini, phylogenetic divergence was marked by a significant negative directional change in size (as well as evolvability, in the case of primates)^12^, suggesting that heterothermy is a derived condition in these orders while homeothermy, or at least the potential for permanent homeothermy, was the ancestral condition in Boreoeutheria. This interpretation is in keeping with earlier studies indicating that the incidence of heterothermy is strongly lineage dependent^14, 21, 139^. Our model thus predicts a loss of microRNA families in heterothermic clades of Boreoeutheria relative to sister clades that are homeothermic, contrary to the overall trend in mammalian evolution^82^.

In order to address this question, we first verified the effect of phylogenetic order on the sizes of 40 primates, including six species in which torpor has been documented^140, 141^: Catarrhini > Platyrrhini ∼ Lemuriformes > Lorisiformes > Heterotherms (*P* = 1.73e-11; Tukey-Kramer) (Supplementary Table S8). Appying MirMachine^107^ to the genomes of 33 of these species, we observed a similar ranking of microRNA families (*P* = 3.64e-18): Catarrhini > Platyrrhini ∼ Lemuriformes > Lorisiformes (Steel-Dwass, nonparametric). Thus, size accounted for 62% of the variation in *miRNA.Fam* (*P* = 3.90e-08). When the mass data within each order were centered on a common scale, body temperature varied negatively with size (*R*^2^ = 0.30, *P* = 4.25e-03, n = 23), provided that heterotherms were excluded. In other words, among primates presumed to be homeothermic, size was positively correlated with the number of microRNA families, secondary to phylogeny, but negatively related to body temperature. Assuming that cellular *MR* is proportional to *miRNA.Fam*, this combination of oppositve trends is indicative of thermoregulation and was lacking among the heterothermic primates. Furthermore, the relative strengths of association of *T*_b_ and *miRNA.Fam* with mass (*R*^2^ = 0.30 and 0.62, respectively) suggest that in response to an evolutionary change in size, body temperatures are less adaptable than cellular metabolic rates^27^.

Analysis of evolutionary rates across a phylogeny of 544 mammals reached the same conclusion^27^: while exhibiting a general downward trend, the branchwise evolution of *T*_b_ was relatively static compared with the rates of change of b*MR* and ambient temperature, consistent with an adaptive barrier to the emergence of higher, but not lower body temperatures. As mammals diversified in size in response to cooler environments^27, 143^, multiple anatomic and physiological mechanisms were available to adjust overall *MR* without altering *T*_b_ or cellular *MR*, including changes in thermal conductance^28, 29^ and relative organ size^144^, particularly that of skeletal muscle^20, 138^. However, shifting to a higher *T*_b_ and cellular *MR* while maintaining the strict relationship between cell size and rate of protein synthesis during the cell cycle would have depended on the ability to synchronize the expression of hundreds of regulatory genes with the activities of their target pathways. Just such a role has been proposed for microRNAs: microRNA activity was essential to buffer the effect of temperature fluctuation on photoreceptor development in Drosophila^145^. When fruit fly larvae were genetically engineered to decrease *MR*, thereby doubling the time to eclosion, the entire microRNA repertoire was rendered dispensable for normal development^146^. According to the mechanism proposed by Cassidy et al (2019)^146^, the effectiveness of transient signaling by a transcription factor depends on the timely decay of its concentration. Since this decay rate is inversely related to the rate of protein synthesis, as metabolic rate increases, greater repressor activity is needed to maintain the same, evolutionarily conserved, rate of decay. By extension, increased repression by microRNAs may have been necessary to maintain the orderly unfolding of developmental programs in warmer-bodied, late-branching mammals.

Conversely, bats are heterotherms that alternate between active thermoregulation to maintain euthermy and torpor, a state in which they are essentially ectotherms^147^. According to the mechanism proposed above, the timing of transient signaling that is microRNA-dependent during euthermy may not be disrupted in the transition to torpor if accompanied by a temperature-related decrease in the rate of Ago2-loading, which is normally rate-limiting for global microRNA activity^148, 149^. The implied ratchet effect, which would make decreases in body temperature more likely than increases, may inform the previously noted trend of birds and mammals to evolve towards cooler body temperatures from warmer-bodied ancestors^27^.

The majority of studies on the evolution of mammalian endothermy have focused on adaptations for the generation, conservation and dissipation of heat^138, 150^, rather than on the mechanism of thermostatic control^26^. Our analysis of the relationships among *cMR*, *miRNA.Fam* and *T*_b_ in mammals has yielded evidence consistent with multivariate selection for stable and efficient thermoregulation. Speakman and Król (2010)<u>^1^</u> have argued that evolutionary fitness must depend more on the realized rate of energy expenditure than on basal metabolic rate (*bMR*), and have proposed that variation in field *MR* (*fMR*) will better reflect the outcome of selection against the risk of hyperthermia. Likewise, this is the underlying assumption for the inference of metabolic rates from daily energy expenditure in models of species niches and their biogeographical distribution^151, 152, 153^. It follows that if variation at the upper end of the range of *cMR*s has been constrained by the ability to dissipate heat, this will be better reflected by the variation of *fMR* with respect to *T*_a_, than by the variation of *bMR* with respect to *T*_b_.

We compared these four parameters in 50 species common to the datasets of Clarke et al (2010)^108^ and Capellini et al (2010)^154^. In the latter, *fMR* had a lower phylogenetic signal than *bMR* in both Metatheria and Eutheria, consistent with a greater role for selection in shaping its distribution (Figure 10). In our derived dataset, the log-based slope of *fMR* (0.5872) was shifted by 0.07 towards greater dependence on surface area than the slope for *bMR* (0.6560) (*P* = 0.0191, ANCOVA), similar to the differences that we found in mammals between the phylogenetic slope of log *rMR versus* log *M* (0.77), on the one hand, and the optimal scaling of *cMR* with respect to *miRNA.Fam* (0.71) on the other, as well as between the optimal *cMR* scaling of early- *versus* late-branching mammals (0.78 versus 0.70) (Supplementary Table S3).

**Figure 10.**
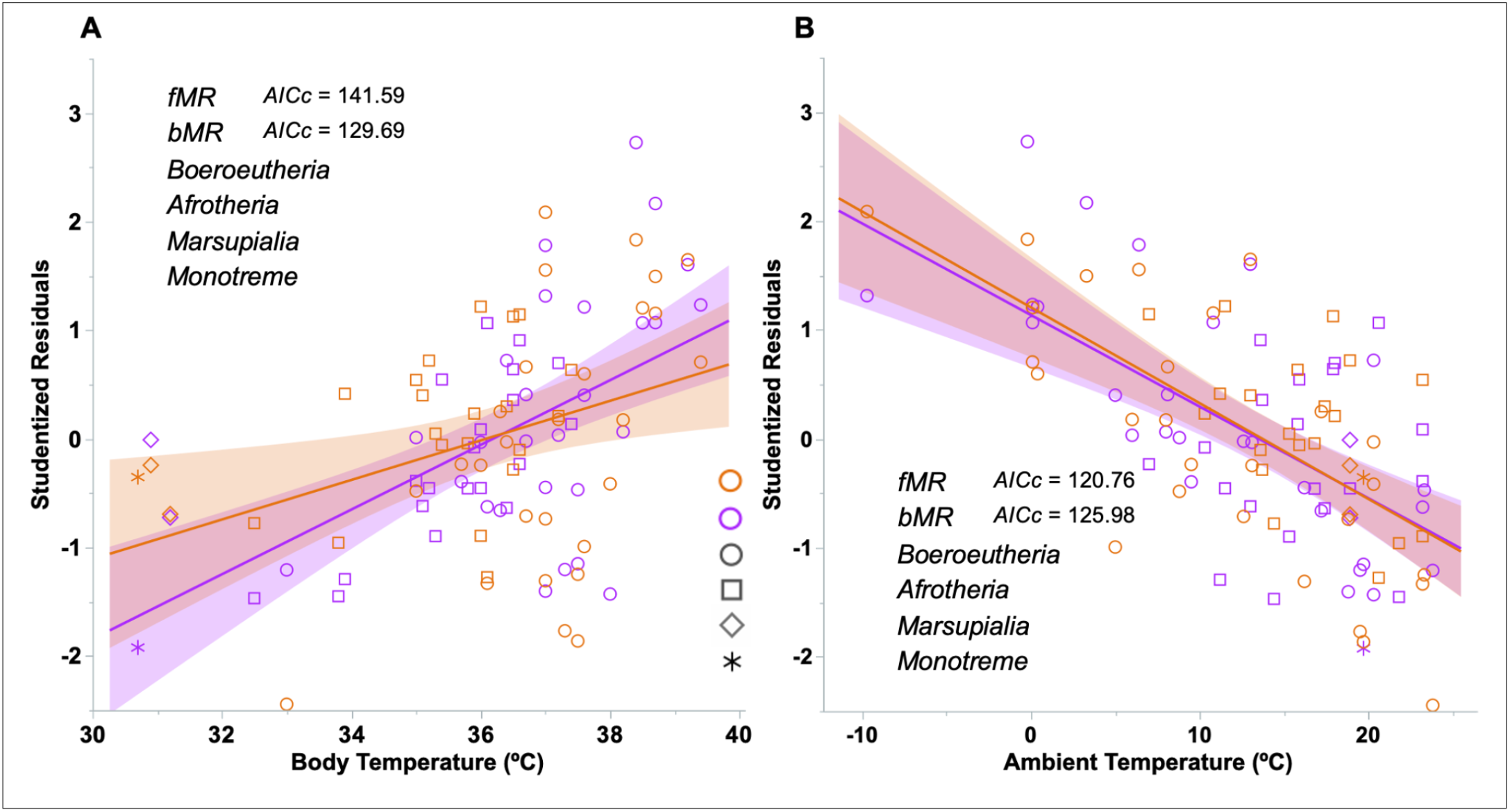
Variation in mass-independent residuals of *rMR* and *fMR* with respect to *T*_b_ and *T*_a_. Species included one Monotreme, two Afrotheria, 20 Marsupials and 27 Boreoeutheria.

As expected, mass-independent residuals from log *bMR* exhibited less variance with respect to body temperature than did the residuals of log *fMR*, but the best predictor for both *bMR* as well as *fMR* was ambient temperature, not body temperature (Figure 10), in agreement with the observations of Avaria-Llautureo et al (2019) in large mammalian trees^27^. These authors found that while the rate of evolution of *bMR* was coupled to *T*_a_ in 75% of all branches, it was coupled to *T*_b_ in only 29% of all branches^27^. In the present study, given the credibility (AICc) of each model and the nearly identical slopes for *fMR* and *bMR* relative to ambient temperature (Figure 10B), we infer the following ranking of significance of the factors that have shaped variation in mammalian metabolic rate: *T*_b_ < *bMR* < *fMR* < *T*_a_. This order of importance appears to reconcile our observations, as well as those of Avaria-Llautureo et al (2019)^27^ with the HDL theory of Speakman and Król (2010)^1^.

There is additional biogeographic and physiological evidence for the notion that the maximal rate of heat dissipation and risk of hyperthermia have exerted a hard constraint on the upper end of variation in body temperature and metabolic rate^155^. Phylogenetically-informed patterns of geographic distribution indicate that the diversification of mammalian morphology has been asymmetric, with sizes increasing in most orders^85^, but evolving more rapidly at higher latitudes and during cooler geologic eras^141^. Similarly, the diversification of the limits of tolerance to cold temperatures evolved faster than the limits of tolerance to heat and fit a biased OU model, *i.e.* stronger selection on the upper critical temperatures^155, 156, 157, 158, 159^. To some extent the asymmetry may reflect a smaller range of habitat variation in warm climatic niches together with more options for behavioral, as opposed to physiological adaptation^159^, but additional evidence points to the operation of a physiological constraint:

i. Thermal performance curves in ectotherms, notably lizards, are markedly asymmetric^151, 152, 160, 161, 162, 163;^
ii. *T*_a_ is a much better predictor than *T*_b_ of the thermodynamic gradient that mammals are able to maintain^158^ (Supplementary Figure S7);
iii. After controlling for mass and phylogeny, the anti-correlation between basal metabolic rates and the upper limit of the thermoneutral zone (the temperature above which cooling mechanisms must be activated in order to prevent hyperthermia) was three times steeper than the anti-correlation with the lower temperature limit (below which cold-induced thermogenesis is activated)^164^;
iv. Relative to unselected lines, inbred lines of bank voles selected for higher *bMR*s under thermoneutral conditions exhibited elevated body temperatures at 34°C, indicative of a diminished capacity for effective thermoregulation and increased risk of hyperthermia^165^.

Our “sliding window” analysis revealed an asymptotic, temperature-related decrease in optimal *cMR* scaling relative to *miRNA.Fam* in later-branching Boreoeutheria. The finding that the optimal allometric slope approached that of field metabolic rate (Figure 9), suggests that the *cMR*-*miRNA.Fam* relationship has been conditioned by physiological limits on the rate of heat dissipation^1^ and implies that as evolving clades of Boreoeutheria increased in size^85, 143^ and/or rate of heat generation^29, 68^, they were forced to negotiate a trade-off between gains in power and thermodynamic efficiency (Δ*T*_b_-*T*_a_), on the one hand, and an increased risk of hyperthermia, on the other. Such a trade-off would be sufficient explanation for the asymmetric distributions of the physiological variables *C*_min_ (thermal conductance) and *U*_CT_ (upper limit of the thermoneutral zone) in Boreoeutheria^157, 158, 159, 166^ (Figure 11).

**Figure 11.**
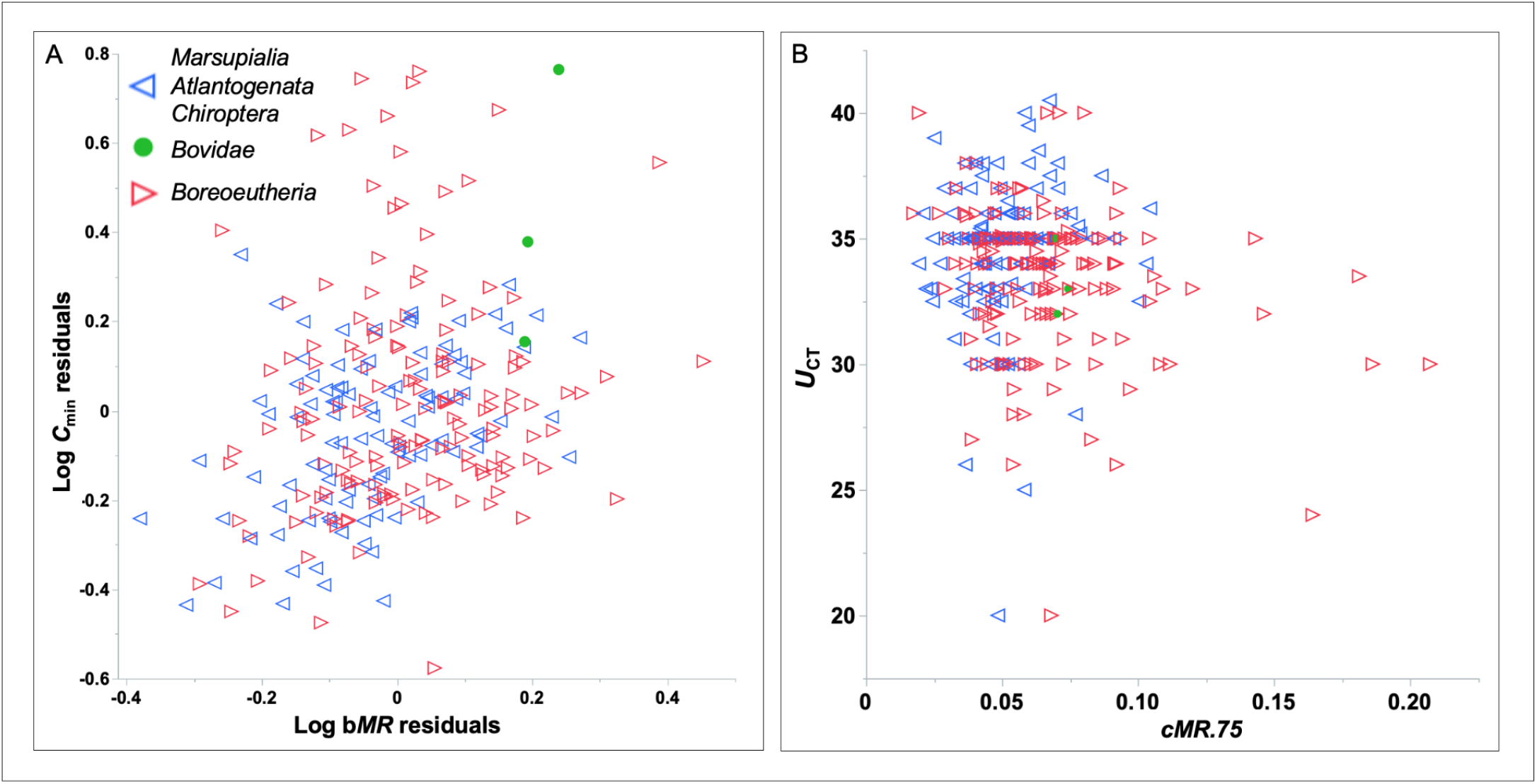
Thermoregulatory Status and the Distributions of Log C_min_ and U_CT_. 276 mammals from the dataset of Khaliq et al (2014)^164^ were divided between Chiroptera and earlier-branching mammals (low incidence of homeothermy) and later-branching Boreoeutheria. Bovids are highlighted because of the well-known constraint that high ambient temperature imposes on the rate of lactation (for references, see Speakman and Król, 2010^1^).

Homeothermic Boreoeutheria exhibited a combination of trends, higher *C*_min_ and lower *U*_CT_, that are consistent with adaptation and constraint at the higher end of the ranges of mammalian *T*_b_ and *cMR*. Later-branching Boreoeutheria predominated in the upper one eighth of the distribution of Log *C*_min_ residuals (*P* = 2.85e-05, □^2^) (Figure 11A). This subset, with 0.75 °C higher *T*_b_’s, and 29% higher *cMR*s than mammals in the lower seven/eighths of the distribution, had *U*_CT_’s that were 2.7°C lower, despite experiencing similar ambient temperatures (Supplementary Table S10).

The adaptive value of repression by individual microRNAs operating in gene regulatory networks has been attributed to their ability to canalize phenotypic expression by buffering the effects of environmental or allelic variation^145, 146, 167^, or to set thresholds for cell fate determination^168, 169^. The ability of each microRNA seed to regulate many targets may have facilitated the evolution of regulatory networks of increasing complexity^170^. Given the potential for microRNA regulation to increase the heritability of adaptive traits^171^, the enlargement of microRNA and target repertoires has been proposed as a key mechanism in the diversification of animal cell types and morphologies^81, 102, 103, 115, 172, 173, 174^. However, relatively little attention has been paid to the nature of global microRNA activity^55, 56, 175, 176, 177^ or its relationship to cellular metabolic rate^146^. Our model predicts that compensatory noise suppression by microRNAs should be proportional to body temperature, since temperature is the primary determinant of the rate of protein synthesis, and therefore of intrinsic stochastic translation noise^46, 178^.

Post-transcriptional regulation of gene expression (“mRNA processing”) is the most constrained gene category in the evolutionary history of placental mammals^179^, which suggests a significant fitness cost for any loss in efficiency of transcript processing as mammals evolved faster and more complex gene regulatory networks.

The present study shifts the focus of research on the evolution of metabolic rate from adaptations of mass-independent factors to the efficiency of the cell mass-dependent rate of protein synthesis. This approach has revealed an unexpected degree of integration of the microRNA-target apparatus into the energy economy of the cell, and reinforces earlier evidence for an energy constraint on the microRNA-dependent ratio of rates of translation of low- and high-abundance transcripts^50^. Taken together, these observations are consistent with selection for co-adaptation of cellular *MR* and global microRNA activity based on metabolic efficiency, thereby facilitating a shift in the Boreoeutherian lineage to higher body temperatures, faster metabolic rates, and permanent homeothermy.

## DATA AVAILABILITY

The original data with accompanying references and the results of all analyses referred to in the text are provided in the Supplementary Information. Phylogenetic tree matrices are available upon request.

## ACKNOWLEDGMENT

BF is supported by the Tromsø forskningsstiftelse (TFS) [20_SG_BF ‘MIRevolution’].

## AUTHOR CONTRIBUTIONS

Conceptualization: BF and TS; Methodology: TS; Investigation: BF and TS; Writing – Original Draft: TS; Writing – Review & Editing: BF and TS; Visualization: BF and TS; Funding Acquisition: BF.

## STAR METHODS

### RESOURCE AVAILABILITY

#### LEAD CONTACT

For further information requests should be directed to Thomas Sorger.

#### MATERIALS AVAILABILITY

This study did not involve any physical material.

#### DATA AND CODE AVAILABILITY

The data compiled for this study are listed in Table S1 in the Supplementary Information. All other data compilations are referenced in the main text.

This paper does not report original code.

### METHOD DETAILS

#### MicroRNA Families, Genes and Targets

The numbers of conserved microRNA families and genes for 20 vertebrate species were obtained from MirGeneDB, a manually curated database of microRNA-encoding genes^67^. Conserved microRNAs were defined as those shared by two or more species. The number of conserved mammalian mRNA targets was extracted from the TargetScan8.0 database^63, 86^ and filtered for targets corresponding only to conserved microRNAs.

#### Other Genomic Variables

Values for genome size and number of coding genes were obtained from the NCBI genome assembly database (as of March, 2022).

#### Body Temperature, Ambient Temperature and Thermal Conductance

For the model presented in Figure 9, body temperatures and ambient temperatures for mammals were obtained from Clarke et al (2010)^108^, excluding multivariate outliers (mass, *bMR*). Mean differences (*T*_b_ - *T*_a_) of 24°C and 16°C were estimated for Boreoeutheria (n = 318) and earlier-branching mammals (n = 94), respectively. A gradient of 20°C for birds < 1,200 g was estimated from mean body temperatures of 40°C (n = 203)^29^ and a mean ambient temperature of 20°C, the average of maximum and minimum temperatures across the entire avian geographical distribution (n = 157)^164^. Upper and lower critical temperatures were drawn from Khaliq et al (2014)^164^ (n = 242) after excluding multivariate outliers (mass, *bMR*). Mass-independent residuals for thermal conductances (*C*_min_) and basal metabolic rates were derived from the dataset of Fristoe et al (2015)^29^, excluding outliers.

### QUANTIFICATION AND STATISTICAL ANALYSIS

#### Body Mass, Resting Metabolic Rates and Temperature Correction

Mass and rates of oxygen consumption and body temperature were drawn primarily from the compilations of White et al (2006)^76^ and Gillooly et al (2017)^77^ (Supplementary Table S1). Where multiple values had been recorded at different temperatures, we used the value nearest to the standard body temperature (see below). The original reference for each species is listed in Table S1 of the Supplementary Information. Resting metabolic rates (*rMR*s) were corrected to 38^°^C for endotherms, and 25^°^C for ectotherms, as *per* White et al (2006)^76^ and Gillooly et al (2017)^77^. Since oxygen consumption data were available only for *X. laevis*, and *X. tropicalis* adults are approximately half the size of adult *X. laevis*, we assumed 0.54 the *rMR* of *X. laevis* (assuming *rMR*_2_/*rMR*_1_ = *M*_2_ ^0.88^/*M*_1_ ^0.88^ for amphibia).

To complement the number of primate microRNA families obtained using MirMachine^107^, mass and uncorrected temperature data were obtained primarily from the AnAge database^180^ and, in a few instances, from the Animal Diversity Web^181^. We used the following formula to center the log_2_*M* data within each order on the same scale: x_cen_ = 50 + 50•(x_i_ - x_avg_)/(x_max_- x_min_).

#### Phylogenies

“Phytools” (Revell, 2012)^182^ was used to generate a 20-species vertebrate tree and phenogram, 14-species tetrapod tree and 10-species mammalian tree, based on divergence times drawn from Álvarez-Carretero et al (2022)^79^ for mammals, and Irisarri et al (2017)^80^ for earlier-branching vertebrates. Trees were made ultrametric and scaled to unit height using Mesquite © (2021)^183^.

In order to model the evolution of *miRNA.Fam*, a matrix of interspecific distances was assembled based on the number of conserved microRNA families (MirGeneDB 2.1)^67^, without considering losses or gains unique to each lineage. A distance tree was inferred by minimal evolution^184^.

#### Statistics

Phylogenetic linear regression was carried out using the *R: CAPER* package^185^ using the maximum likelihood estimate between 0.0-1.0 for Pagel’s *lambda* parameter^6^. All *R*^2^ values were adjusted for the number of predictors.

Multivariate correlations, screening for outliers, ANCOVA and ordinal logistic regression were performed in JMP®16. Cows were considered outliers with respect to mass and, as in the case of all ruminants, metabolic rate (White and Seymour, 2005)^186^. All other statistical tests were performed in R (2020)^187^.

#### OU versus BM comparison and simulation

BM and OU models of the variation of *cMR* with respect to number of microRNA families were evaluated with the package mvSLOUCH^8^, with six repetitions of the hill-climbing algorithm for the OU process using two adaptation matrices (Diagonal and DecomposableReal). Preliminary validation based on the AICc criterion revealed that for the 14 tetrapod model, OU processes were misclassified as BM for 50% of the simulations, while 10% of the BM simulations were misclassified as OU. The corresponding rates of misclassification for the 10-mammal model were 75% and 5% (n = 40, Supplementary Table S3). The rates of correct classification for the BM and OU simulations were equalized by adjusting the AICc scores +/- 2.1 for the tetrapod model, or +/- 4.0 for the mammalian model (Supplementary Table S4).

The Phylogenetic EM package^10^ was used to simulate *cMR* and *miRNA.Fam* distributions using variance/covariance matrix for *cMR.75* on *miRNA.Fam* as the selection strength matrix, with the median setting (3) for selection strength (Supplementary Figure S4). The average values for all mammals *cMR.75* and *miRNA.Fam* were assigned to the first adaptive shift (divergence of mammals), and the averages for all placentals were assigned to the second adaptive shift (divergence of Atlantogenata or Boreoeutheria or Euarchontoglires). Ten simulations were generated and averaged for each shift or pair of shifts.

#### Bayesian inference of the branch-wise rates of evolution

To generate variable rate trees for each trait, the VarRates algorithm of *BayesTraits*^188^ (version 4, www.evolution.rdg.ac.uk) was run for 12.5 million iterations with 20% burn-in and sampled every 1000 trees, yielding 10,000 trees fit to the distribution of either log_2_*M*, *cMR.75*, *miRNA.Fam* or *T*_b_ ^7^. Bayes factors were obtained by comparing median likelihoods of the 100 trees with the highest log likelihoods and the 100 trees with the smallest non-zero sum of scale changes (constant rate model). The latter was equivalent to the median log likelihood of random walk models for the reference tree (constant rate model). For the temperature data and reference tree, for example, the median log likelihood of 10,000 random walk models was −25.90, while the median log likelihood of the 100/10,000 variable rate trees with the smallest non-zero sum of scale changes was −26.62.

For each trait, correlations among the sets of branch lengths between the divergence of Theria and the divergence of Primates were obtained from the median branch lengths of the best 100 (1%) variable rate trees.

#### Variable Rate Regression and Inference of Internal Node Values

In order to evaluate the dependence of *cMR.75* on *miRNA.Fam*, while controlling for size (log mass) and body temperature, Bayesian regressions were run for 2.25 million iterations with 20% burn-in and sampled every 1000 trees (BayesTraits). Model statistics represent median values for the models with the 5% highest likelihood (100 trees).

Internal node values (BayesTraits) for log_2_ *M*, *cMR.75*, *T*_b_ and *miRNA.Fam* were inferred from the random variation model run for 8.0 million iterations with 20% burnin and sampled every 1,000 trees.

#### Quantification of Model Interactions between Reference microRNAs and their 3’-UTR Target Sites

The entries in the TargetScan database “Conserved Site Context Scores”^63, 86^ were filtered for the reference microRNAs listed for each species in MirGeneDB^67^. For each distribution of CWCS scores, the median value was used to estimate sensitivity to repression. In each species, a non-redundant list of target genes was used to compute the average number of 3’-UTR target sites/gene and the length distribution of the corresponding target proteins between 500 - 5,000 amino acids. The relative ‘AA cost’ was obtained from the weighted sum of nine 500 AA-length bins between 500 - 5,000 AA (Supplementary Tables S2).

