## Supplementary Information for "Rapid Adaptation of Cellular Metabolic Rate to the MicroRNA Complements of Mammals and its Relevance to the Evolution of Endothermy"

**Table S1**. Summary of phenotype data for all variables for 20 vertebrate species in MirGeneDB 2.1[1] (abbreviations as in main text). MicroRNA numbers do not include novel genes *i.e.* genes unique to a single species. Rat and rabbit data, and the second set of data for chicken, were used for the comparison of specific rates of protein synthesis. [a]Values were adjusted by subtracting 25 genes/families unique to mouse and rat. These are the source data for main text Figures 1 - 7, Supplementary Figures S2 - S6 and Supplementary Tables S3 - S7.

| **Species** | ***miRNA. Fam*** | ***miRNA. Gen*** | $T_b$ (°K) | ***Gnm*** (x100 Mb) | ***Cdg*** (x 1000) | ***rMR*** (ml $O_2$/h) | ***Mass*** (g) | **Ref's.** |
|---|---|---|---|---|---|---|---|---|
| Human | 265 | 563 | 309.95 | 31.00 | 17.04 | 13584 | 60500 | 2 |
| Monkey | 217 | 502 | 309.95 | 29.71 | 21.07 | 2358 | 6230 | 2 |
| Mouse | 199[a] | 427[a] | 309.95 | 27.28 | 22.34 | 50.94 | 31.33 | 3 |
| Guinea Pig | 184 | 402 | 312.15 | 27.23 | 19.98 | 322.8 | 629 | 4 |
| Cow | 212 | 457 | 311.85 | 27.16 | 20.77 | 53441 | 347000 | 4 |
| Dog | 189 | 433 | 311.55 | 23.97 | 19.74 | 7160.4 | 30000 | 5 |
| Armadillo | 169 | 380 | 307.65 | 36.32 | 22.03 | 1102.8 | 3510 | 4 |
| Tenrec | 160 | 344 | 305.15 | 29.47 | 20.00 | 97.1 | 175.8 | 6, 7 |
| Tasmanian Devil | 145 | 396 | 308.95 | 30.87 | 19.99 | 1543 | 5775 | 4 |
| Opossum | 167 | 500 | 305.75 | 35.98 | 20.65 | 87.24 | 104 | 4 |
| Platypus | 136 | 388 | 307.15 | 18.59 | 17.84 | 255.98 | 693 | 4 |
| Zebra Finch | 115 | 257 | 313.95 | 10.56 | 16.52 | 33.07 | 15.00 | 8 |
| Pigeon | 121 | 257 | 313.95 | 11.09 | 15.60 | 289.1 | 414 | 9 |
| Chicken | 136 | 286 | 313.25 | 10.53 | 17.48 | 929.8 | 2710 | 4 |
| Alligator | 113 | 283 | 308.15 | 21.62 | 18.96 | 47.17 | 1287 | 4 |
| Turtle | 123 | 300 | 298.15 | 23.66 | 21.19 | 6.614 | 219.00 | 10 |
| Lizard | 118 | 267 | 303.15 | 17.99 | 19.24 | 0.671 | 4.50 | 4 |
| Frog | 95 | 343 | 298.15 | 14.51 | 21.90 | 3.985 | 31.90 | 4 |
| Zebrafish | 105 | 406 | 301.15 | 13.73 | 27.46 | 0.009 | 0.10 | 11 |
| Catshark | 87 | 244 | 285.15 | 44.71 | 28.05 | 29.550 | 500 | 12 |
| Rat | 164[a] | 395[a] | 311.15 | 26.48 | 23.35 | 240.0 | 250.0 | 13 |
| Rabbit | 184 | 389 | 311.95 | 27.37 | 20.55 | 1368.0 | 2150.0 | 14 |
| Chicken | 136 | 286 | 313.25 | 9.53 | 16.88 | 1253.78 | 1493.0 | 15 |

**Tables S2.** Analysis of TargetScan Database for Conserved 3'-UTR microRNA Target Sites.

**A.**Conserved target sites 16[16], [17] were filtered according to the reference microRNA complements of the eight species included in the present study[1]. See main text Figure 8.
Repression/Site: cumulative repression ($C_n$) computed from the median value of the "cumulative weighted context++ scores". AA Cost: the weighted sum of the distribution of polypeptide lengths between 500 – 5,000 amino acids.
∆AA Cost: Repression x AA cost. **B.**The same data standardized according the values in the opossum. [illegible]

**A.**

| Species | Conserv.DB sites | MGDB targets | Sites/Target | Targets/Cdg | Sites/Cdg | Repression /Site ($C_n$) | ∆ Translation | AACost | ∆ AA Cost |
|---|---|---|---|---|---|---|---|---|---|
| Frog | 21,636 | 15,956 | 8.60 | 0.1152 | 0.9913 | 0.1355 | 0.134 | 1385 | 186 |
| Chicken | 47,150 | 36,220 | 10.60 | 0.2546 | 2.6978 | 0.1270 | 0.343 | 1168 | 400 |
| Opossum | 78,080 | 71,274 | 10.71 | 0.3487 | 3.7353 | 0.1118 | 0.418 | 1152 | 481 |
| Dog | 156,369 | 122,139 | 13.70 | 0.5680 | 7.7846 | 0.0994 | 0.774 | 1120 | 866 |
| Cow | 173,993 | 141,928 | 15.22 | 0.5450 | 8.2933 | 0.1179 | 0.978 | 1128 | 1103 |
| Mouse | 195,305 | 118,130 | 17.38 | 0.5065 | 8.8007 | 0.0981 | 0.864 | 1072 | 926 |
| Monkey | 202,546 | 160,091 | 16.62 | 0.5770 | 9.5898 | 0.0981 | 0.941 | 1151 | 1083 |
| Human | 264,563 | 173,422 | 20.24 | 0.6509 | 13.1754 | 0.0931 | 1.227 | 1056 | 1295 |

**B.**

| Species | Conserv.DB sites | MGDB targets | Sites/Target | Targets/Cdg | Sites/Cdg | Repression /Site ($C_n$) | ∆ Translation | AACost | ∆ AA Cost |
|---|---|---|---|---|---|---|---|---|---|
| Frog | 0.277 | 0.224 | 0.803 | 0.330 | 0.265 | 1.212 | 0.322 | 1.202 | 0.387 |
| Chicken | 0.604 | 0.508 | 0.990 | 0.730 | 0.722 | 1.136 | 0.821 | 1.014 | 0.832 |
| Opossum | 1.000 | 1.000 | 1.000 | 1.000 | 1.000 | 1.000 | 1.000 | 1.000 | 1.000 |
| Dog | 2.003 | 1.714 | 1.280 | 1.629 | 2.084 | 0.889 | 1.852 | 0.972 | 1.801 |
| Cow | 2.228 | 1.991 | 1.421 | 1.563 | 2.220 | 1.055 | 2.342 | 0.979 | 2.293 |
| Mouse | 2.501 | 1.657 | 1.622 | 1.453 | 2.356 | 0.878 | 2.068 | 0.931 | 1.925 |
| Monkey | 2.594 | 2.246 | 1.552 | 1.655 | 2.567 | 0.878 | 2.253 | 0.999 | 2.252 |
| Human | 3.388 | 2.433 | 1.890 | 1.867 | 3.527 | 0.833 | 2.938 | 0.917 | 2.693 |

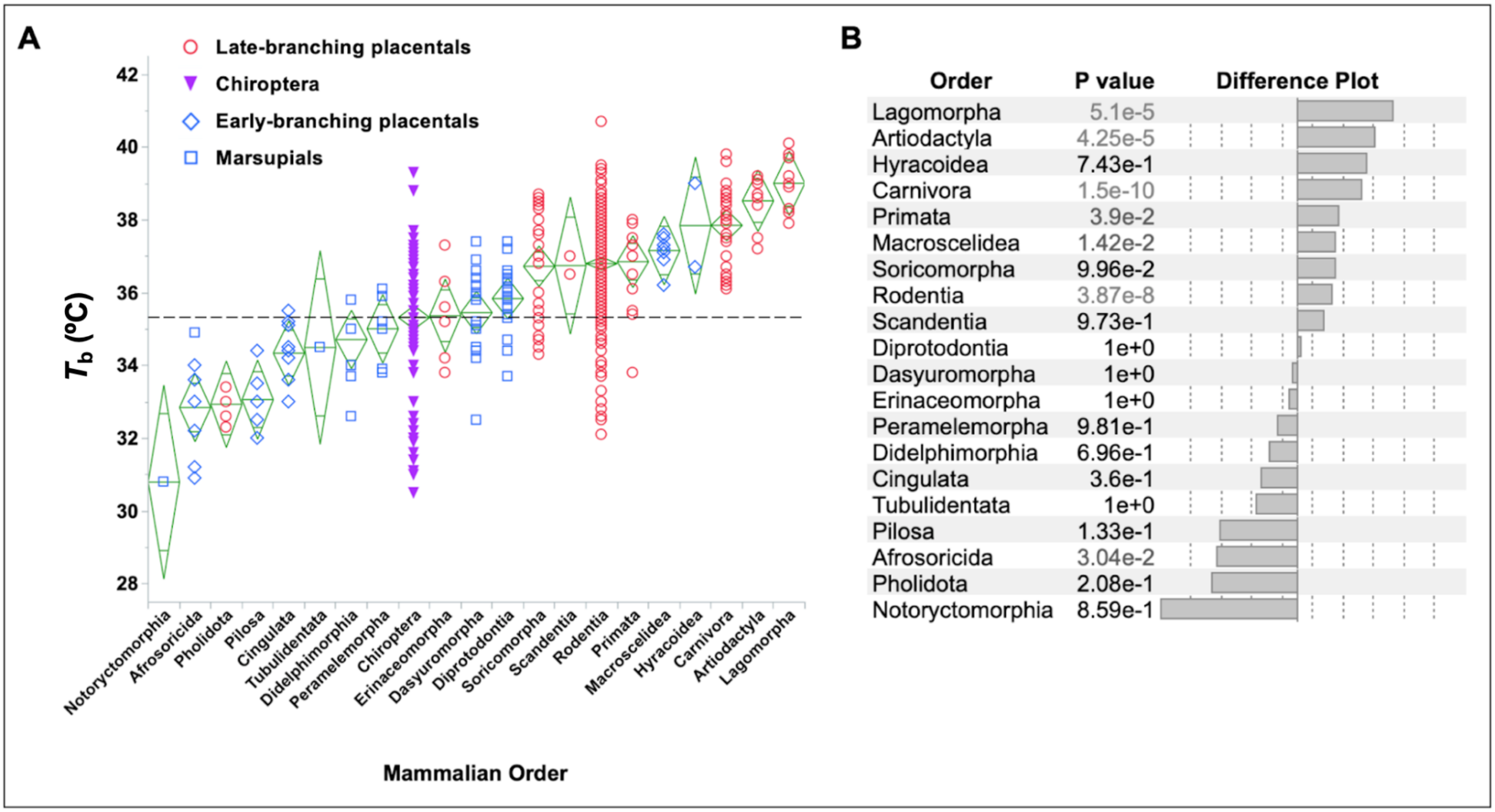

**Figure S1**. Ranking by temperature ($T_b$) of 20 orders of terrestrial mammals.
A. Distribution of $T_b$ by order and species. Diamonds represent means and confidence intervals. The horizontal line represents the mean temperature of Chiroptera (35.3°C).
B. Difference plot. *P* values refer to comparison of each order with Chiroptera (Steel's nonparametric method). Data (n = 412) were drawn from Clarke, Rothery and Isaac (2010)[18].

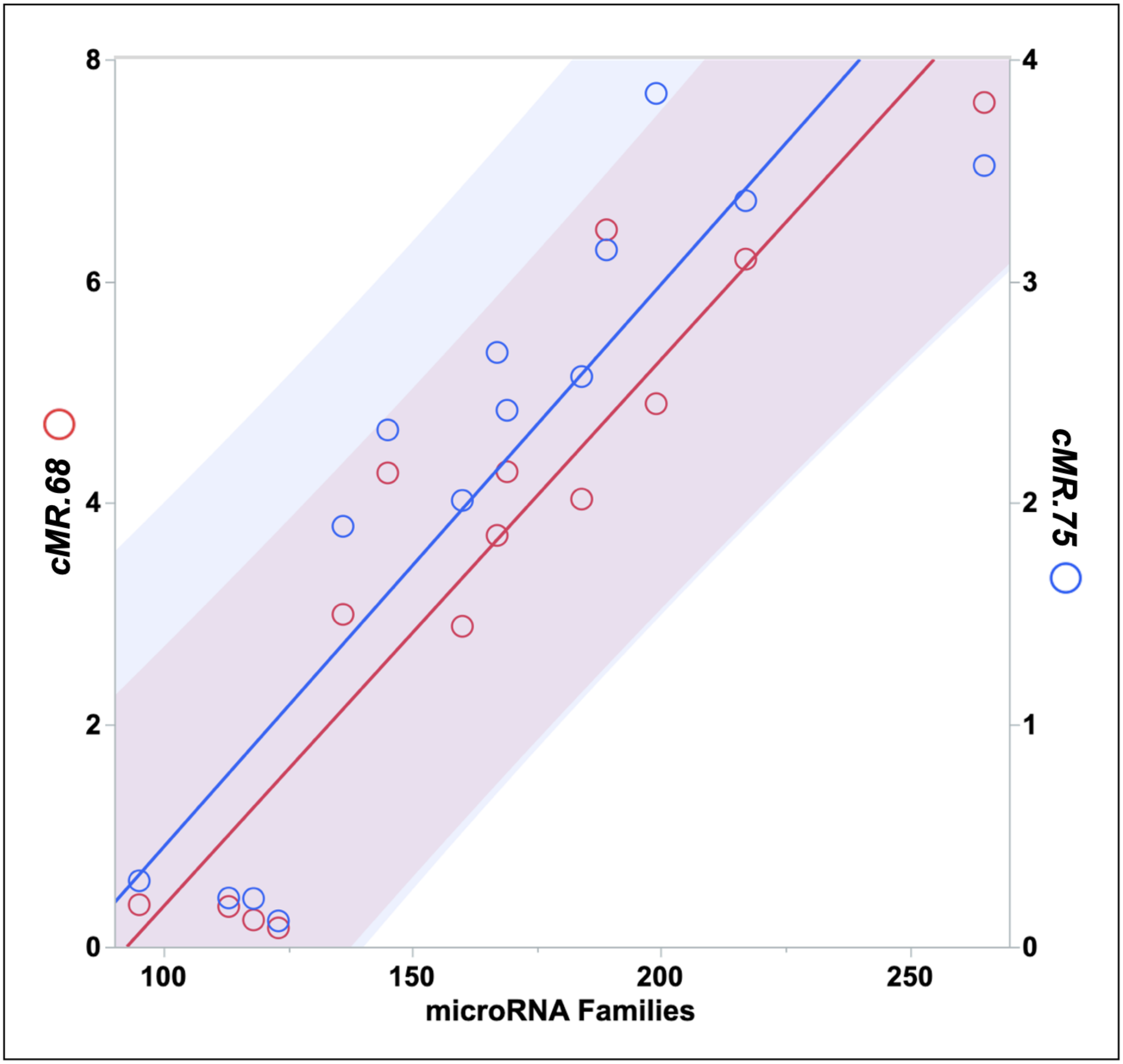

**Figure S2**. Maximum likelihood fitting of *rMR/M*$^{0.68}$ or *rMR/M*$^{0.75}$ to *miRNA.Fam* across 14 tetrapods yields variation that is homoscedastic.
Shaded bands represent the 95% confidence limits. Compare with main text Figure 4A.

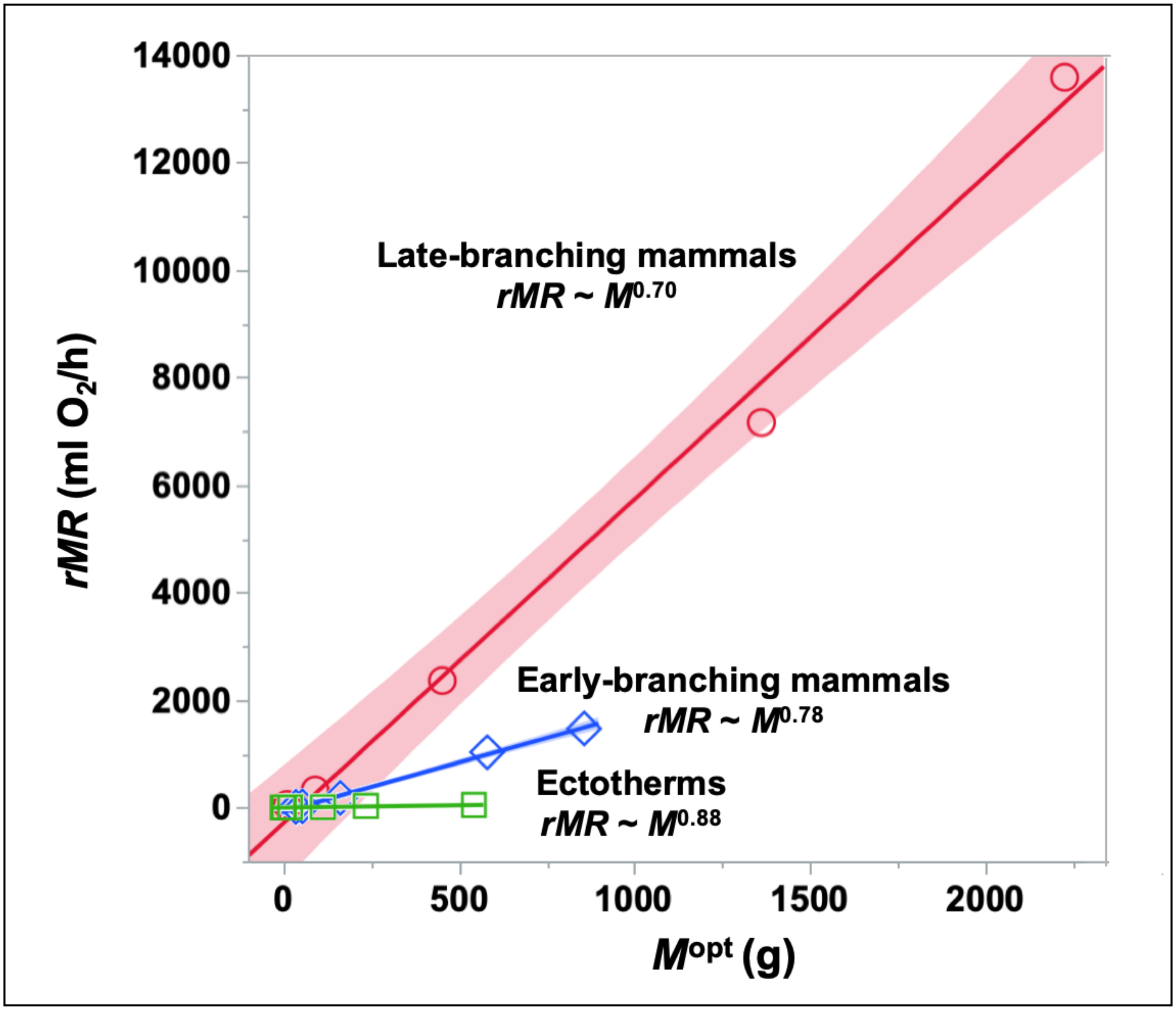

**Figure S3.** Comparison of the linear fit and confidence limits for *cMR*s with slopes scaled optimally with respect to *miRNA.Fam* in five early- and five late-branching mammals, as well as the conventional log-based slope for six ectotherms (refers to main Figure 4A and Supplementary Table S3).

**Table S3**. Summary of Phylogenetic Regressions (constant rate assumption).
Tetrapods included 5 late-branching mammals, 5 early-branching mammals, 3 non-avian reptiles and an amphibian. *cMR* = optimally scaled cellular metabolic rate (*rMR*/$M^{exp}$), *Gnm* = genome size, *Cdg* = number of coding genes. In each case, the dependent variable is written first. These data reference main text Figure 4A.

| Clade | Model | Allometric Exponent | lambda | $R^2$ | P | AICc |
|---|---|---|---|---|---|---|
| **Tetrapods** | **log rMR ~ log M** | $M^{0.78}$ | 1.000 | 0.9443 | 4.28e-09 | 6.30 |
| *n = 14* | **cMR~miRNA.Fam** | $M^{0.68}$ | 0.000 | 0.8427 | 2.26e-06 | 41.92 |
| | **cMR~miRNA.Gen** | $M^{0.67}$ | 0.000 | 0.7593 | 3.02e-05 | 50.63 |
| **Mammals** | **log rMR ~ log M** | $M^{0.77}$ | 0.000 | 0.9832 | 1.36e-08 | -12.65 |
| *n = 10* | **cMR~miRNA.Fam** | $M^{0.71}$ | 0.000 | 0.7777 | 4.55e-04 | 17.40 |
| | **cMR~mRNA.Gen** | $M^{0.72}$ | 0.617 | 0.6120 | 4.60e-03 | 21.65 |
| **Late-branching** | **log rMR ~ log M** | $M^{0.74}$ | 0.000 | 0.9943 | 1.19e-04 | -4.18 |
| *n = 5* | **cMR~miRNA.Fam** | $M^{0.70}$ | 0.000 | 0.5170 | 0.1052 | 17.48 |
| | **cMR~miRNA.Gen** | $M^{0.71}$ | 0.000 | 0.6633 | 5.86e-02 | 13.74 |
| **Early-branching** | **log rMR ~ log M** | $M^{0.75}$ | 0.000 | 0.9853 | 4.91e-04 | -4.91 |
| *n = 5* | **cMR~miRNA.Fam** | $M^{0.78}$ | 0.000 | 0.3352 | 0.1808 | 7.22 |
| | **cMR~miRNA.Gen** | $M^{0.78}$ | 1.000 | 0.8498 | 1.66e-02 | 3.40 |

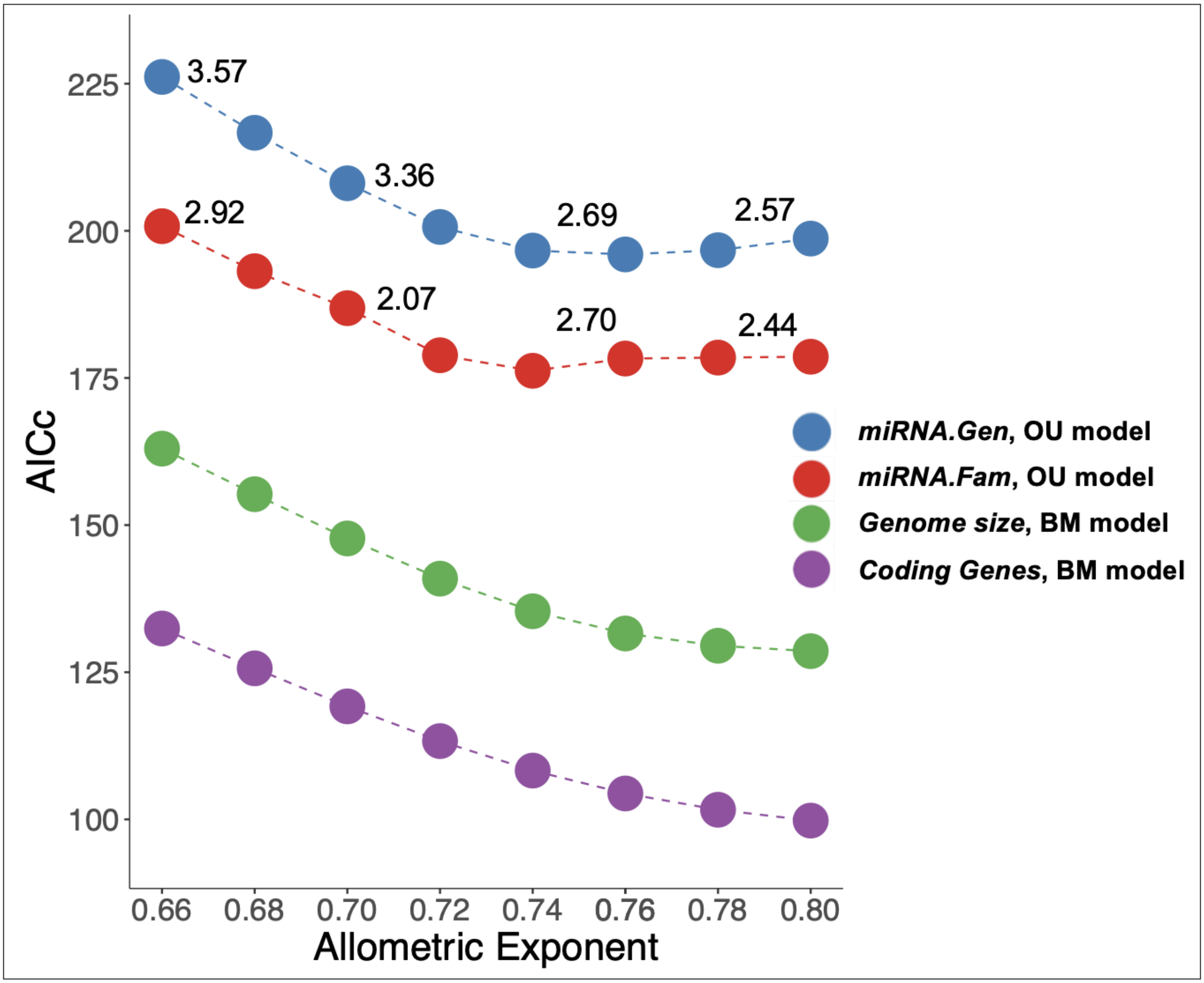

**Figure S4**. Comparison of BM and OU models for the dependence of *cMR* on the number of microRNA families (*miRNA.Fam*), microRNA genes (*miRNA.Gen*), genome size (*Gnm*) and the number of coding genes (*Cdg*) across 14 tetrapods. AICc values were not corrected for bias (see Methods). Values above the *miRNA.Gen* and *miRNA.Fam* data represent uncorrected differences in ∆AICc (BM model - OU model). Compare with main text Figure 5.

**Table S4**. Empirical Correction of Bias for Classification of BM *versus* OU Models.
40 simulated pairs of data (*cMR75*, *mirFam*) were generated for 14 tetrapods or 10 mammals (see Methods). Each simulation was evaluated by the mvSLOUCH algorithm and classified according to the lowest AICc value. These corrections were applied to the AICc values presented in main text Figure 5.

| ***Clade*** | ***Simulations N = 40*** | ***Classification before correction*** | | | ***Classification after correction*** | |
|---|---|---|---|---|---|---|
| | | **OU** | **BM** | ***Correction*** | **OU** | **BM** |
| ***Tetrapods*** n = 14 | **OU** | 20 | 20 | -2.1 | 28 | 12 |
| | **BM** | 4 | 36 | +2.1 | 28 | 12 |
| | | **OU** | **BM** | ***Correction*** | **OU** | **BM** |
| ***Mammals*** n = 10 | **OU** | 10 | 30 | -4.0 | 20 | 20 |
| | **BM** | 38 | 2 | +4.0 | 20 | 20 |

**Table S5.** Branch Length Correlations among the Variable Rate Trees for $Log_2 M$, *cMR.75* and *miRNA.Fam*. (refers to main text Figure 7).
Correlations (lower left) and correlation probabilities (upper right) are shown for the median branch lengths between the divergence of Theria and the divergence of Primates for the 1% most likely trees for each trait.

| **Traits** | **$Log_2$ *M*** | **$T_b$ (ºC)** | ***sMR.75*** | ***miRNA.Fam*** | **PROBABILITIES** |
|---|---|---|---|---|---|
| **$Log_2$ *M*** | | 5.91e-02 | 5.89e-02 | 5.74e-02 | **$Log_2$ *M*** |
| **$T_b$ (ºC)** | 0.7944 | | 6.36e-10 | 2.46e-07 | **$T_b$ (ºC)** |
| ***sMR.75*** | 0.7947 | 1.0000 | | 8.91e-08 | ***sMR.75*** |
| ***miRNA.Fam*** | 0.7974 | 0.9996 | 0.9998 | | ***miRNA.Fam*** |
| **CORRELATIONS** | **$Log_2$ *M*** | **$T_b$ (ºC)** | ***sMR.75*** | ***miRNA.Fam*** | |

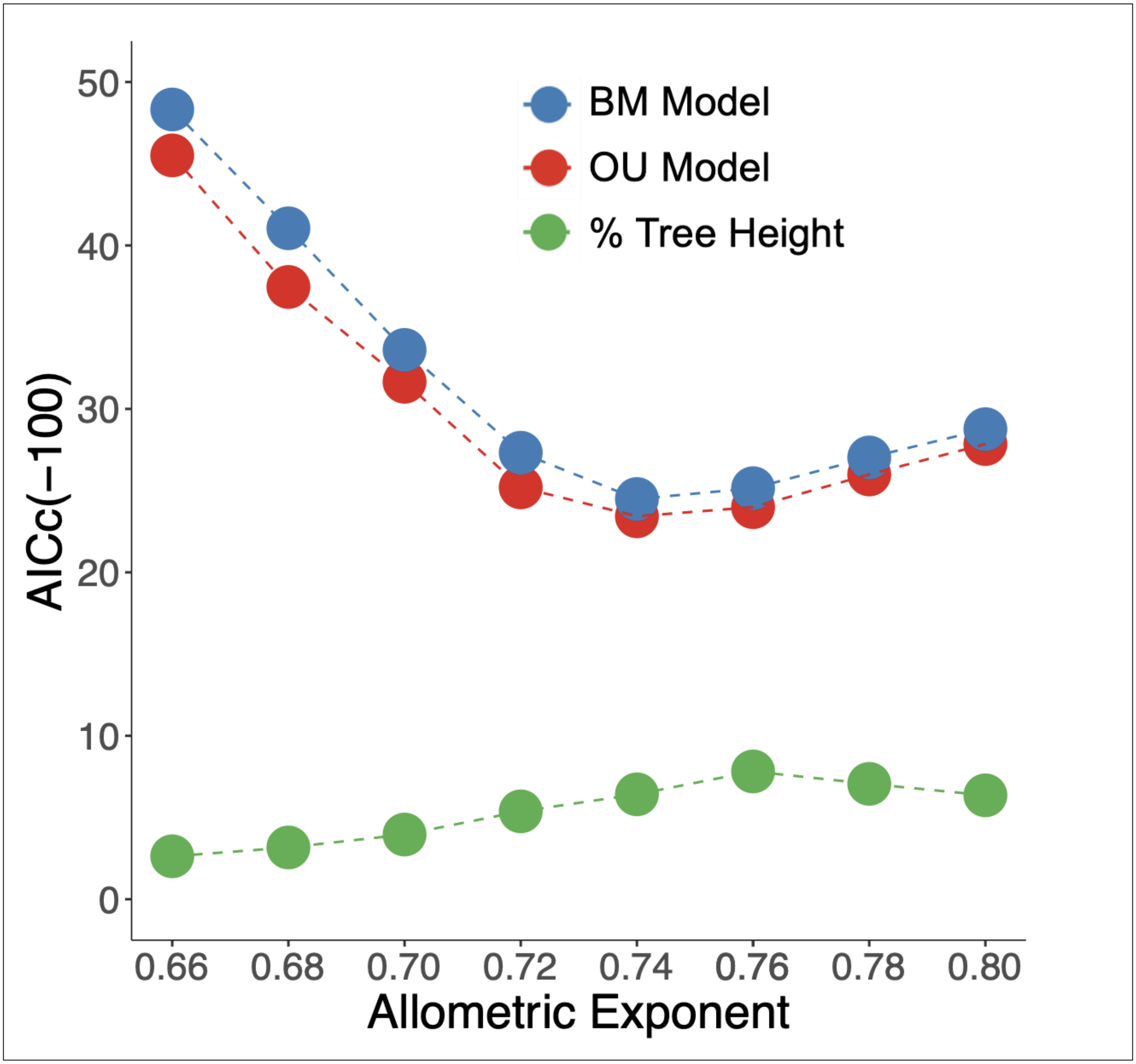

**Figure S5**. Comparison of BM and OU models for the dependence of *cMR* on the number of microRNA families (*miRNA.Fam*) in 10 tetrapods.
Models were based on the variable rate tree for *sMR.75*. ∆AICc (BM-OU) was -0.53 before, and +1.47 after the correction of bias (+/- 1.0) (see Methods). Refers to main text Figure 5.

**Table S6**. ANCOVA and Bayesian Regressions: variable *versus* constant rate models.

Bayes factors (*B*) were obtained by comparing the marginal likelihoods of regressions obtained for models with 10 mammals (excluding cow)[19]. "% neg" refers to the percent of all models in which the coefficient for the second dependent variable was negative, and therefore less likely to be supported. These data provide statistical support for the interpretation of the variable rate trees in main text Figure 7.

| **ANCOVA MODELS** | ***cMR.75 ~ miRNA.Fam*** | ***cMR.75 ~ miRNA.Fam* • $\log_2 M$** | ***cMR.75 ~ miRNA.Fam* • $T_b$** |
|---|---|---|---|
| **Uncorrected/Corrected (•)** | $R^2 = 0.67$; $P = 0.0040$ | $R^2 = 0.88$; $P = 0.0267$ | $R^2 = 0.78$; $P = 0.2133$ |
| **BAYES REGR'N MODELS** | ***cMR.75 ~ $\beta_1$ miRNA.Fam*** | ***cMR.75 ~ $\beta_1$ miRNA.Fam + $\beta_2$ $\log_2 M$*** | ***cMR.75 ~ $\beta_1$ miRNA.Fam + $\beta_2$ $T_b$*** |
| **Variable Rate $R^2$** | 0.8254 | 0.8183 | 0.8329 |
| **Constant Rate $R^2$** | 0.5483 | 0.6790 | 0.5559 |
| **Variable Rate Variance** | 0.0003 | 0.0003 | 0.0003 |
| **Constant Rate Variance** | 0.0017 | 0.0012 | 0.0017 |
| ***miRNA.Fam* Coefficient** | 0.0153 | 0.0140 | 0.0144 |
| **$\log_2 M$ or $T_b$ Coefficient (% neg)** | | 0.0003 (93.8) | 0.0247 (34.6) |
| **Marg. Lh: Variable** | -18.8426 | -27.7445 | -23.0002 |
| **Marg. Lh: Constant** | -20.7936 | -28.0166 | -27.6897 |
| **B Factor** | 3.9020 | 0.5443 | 9.3791 |

**Table S7**. 146 microRNA families in MirGeneDB common to the six Boreoeutheria in this study, including those missing from the genomes of armadillo (red) or tenrec (blue).

Only a single microRNAfamily, MIR-370, is absent from the genomes of both Atlantogenata. MIR-105, MIR-370 and MIR-6715 do not have conserved targets in these six species. These data inform the model represented in main text Figure 8.

| | | | | | | | |
|---|---|---|---|---|---|---|---|
| LET-7 | MIR-1 | MIR-7 | MIR-8 | MIR-9 | MIR-10 | MIR-15 | MIR-17 |
| MIR-19 | MIR-21 | MIR-22 | MIR-23 | MIR-24 | MIR-26 | MIR-27 | MIR-28 |
| MIR-29 | MIR-30 | MIR-31 | MIR-32 | MIR-33 | MIR-34 | MIR-92 | MIR-95 |
| MIR-96 | MIR-101 | MIR-103 | **MIR-105** | MIR-122 | MIR-124 | MIR-126 | MIR-127 |
| MIR-128 | MIR-129 | MIR-130 | MIR-132 | MIR-133 | **MIR-134** | MIR-135 | MIR-136 |
| MIR-138 | MIR-139 | MIR-140 | MIR-142 | MIR-143 | MIR-144 | MIR-145 | MIR-146 |
| **MIR-147** | MIR-148 | MIR-149 | MIR-150 | MIR-153 | MIR-154 | MIR-155 | MIR-181 |
| MIR-184 | MIR-185 | MIR-186 | MIR-187 | MIR-188 | MIR-190 | MIR-191 | MIR-192 |
| MIR-193 | MIR-194 | MIR-196 | MIR-199 | MIR-202 | MIR-203 | MIR-204 | MIR-205 |
| MIR-208 | MIR-210 | MIR-214 | MIR-216 | MIR-217 | MIR-218 | MIR-219 | MIR-221 |
| MIR-223 | **MIR-224** | **MIR-296** | MIR-320 | MIR-324 | MIR-326 | MIR-328 | MIR-330 |
| MIR-331 | MIR-335 | MIR-338 | MIR-339 | MIR-340 | MIR-342 | MIR-345 | **MIR-346** |
| MIR-361 | MIR-362 | ***MIR-370*** | **MIR-374** | MIR-375 | MIR-376 | MIR-378 | MIR-383 |
| MIR-423 | MIR-425 | MIR-430 | **MIR-431** | MIR-433 | **MIR-448** | **MIR-450** | MIR-451 |
| **MIR-452** | MIR-455 | MIR-459 | MIR-483 | MIR-486 | MIR-490 | **MIR-491** | MIR-493 |
| MIR-499 | MIR-504 | MIR-505 | MIR-506 | MIR-542 | MIR-551 | MIR-574 | **MIR-582** |
| MIR-592 | **MIR-615** | MIR-652 | **MIR-653** | MIR-671 | MIR-744 | MIR-873 | MIR-874 |
| MIR-1247 | MIR-1249 | MIR-1251 | MIR-1271 | **MIR-1298** | MIR-1306 | **MIR-1912** | MIR-3059 |
| MIR-3085 | **MIR-6715** | | | | | | |

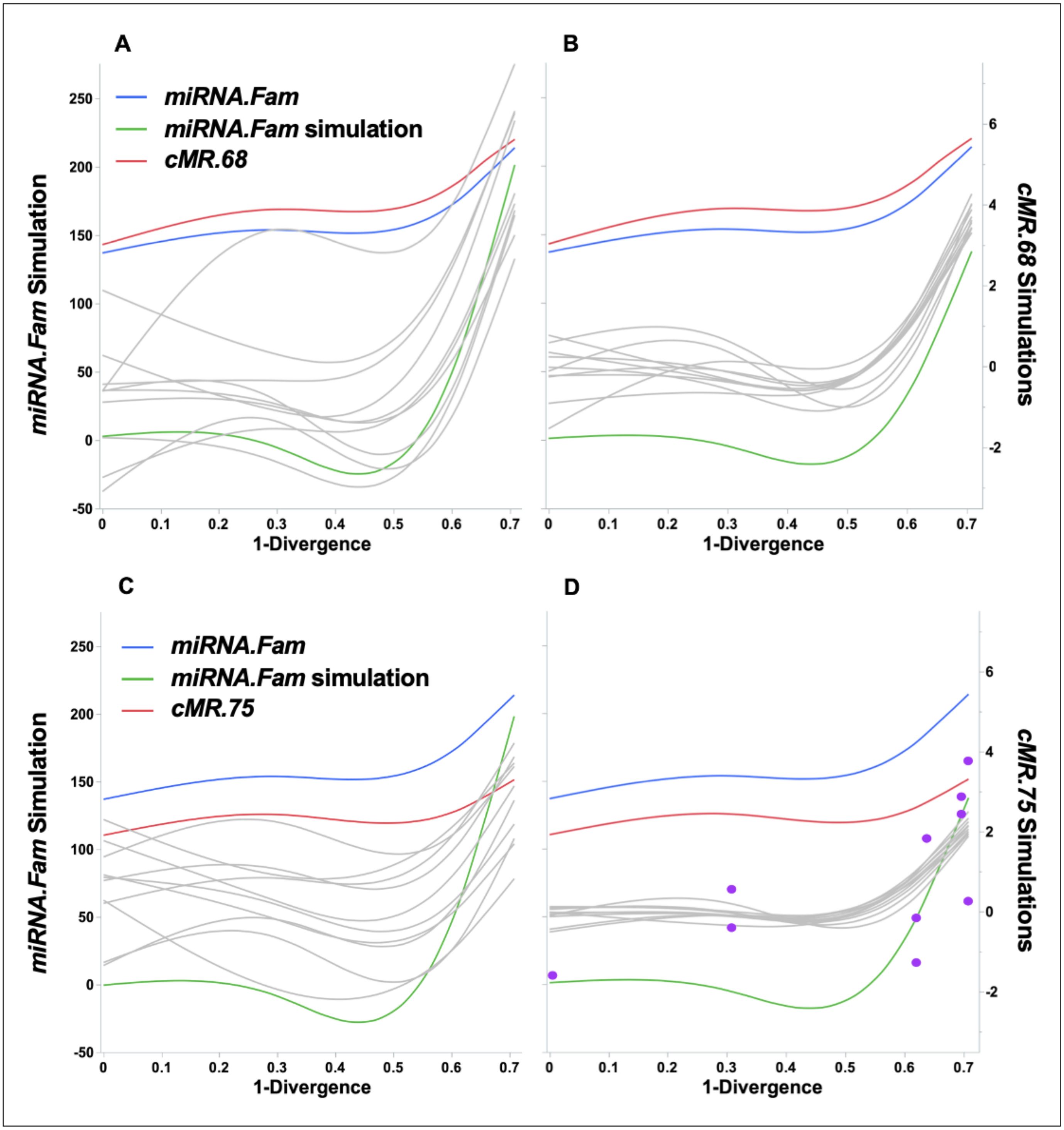

**Figure S6.** Evolution of *miRNA.Fam*, *cMR*.68 and *cMR*.75 in mammals.
Simulations were based on the variance/covariance matrices of *cMR*.68 (A, B) or *cMR*.75 (C, D) with respect to *mirFam*, with an adaptive shift at the divergence of Boreoeutheria (see main text Figure 6).
BM models: A, C. OU models: B, D. The points in panel D (OU) represent the rescaled values of *cMR.75* ($rMR/M^{0.75}$).

**Table S8**. Primate microRNA families detected by MirMachine and their correlated traits.
Reference genomes (NCBI) were annotated by MirMachine (Umu et al, 2022)[20] Trait data were drawn from the AnAgeref database and the Animal Diversity Webref (see Methods). For five pairs of species (asterisks/shaded) the species for which body temperature was available was assigned the number of microRNA families found in another species in the same genus. The following formula was used to center the $\log_2 M$ data within each order on a common scale: $\mathbf{x_{cen} = 50 + 50 \cdot (x_i - x_{avg})/(x_{max} - x_{min})}$.

| ParvOrder | Species | Heterothermy | *miRNA.Fam* | $T_b$ (°C) | Mass (g) | $Log_2M$ | $Log_2M_{cen}$ |
|---|---|---|---|---|---|---|---|
| Catarrhini | *Cercopithecus mitis* | No | 200 | 37.5 | 8648.7 | 13.078 | 41.306 |
| Catarrhini | *Chlorocebus aethiops* | No | 199 | 36.8 | 5620 | 12.456 | 35.695 |
| Catarrhini | *Colobus guereza* | No | 198 | 37 | 10624 | 13.375 | 43.985 |
| Catarrhini | *Erythrocebus patas* | No | 201 | 39.3 | 3000 | 11.551 | 27.531 |
| Catarrhini | *Macaca fascicularis* | No | 207 | 37.6 | 6363 | 12.635 | 37.309 |
| Catarrhini | *Macaca mulatta* | No | 208 | 37.3 | 8235 | 13.008 | 40.674 |
| Catarrhini | *Papio anubis* | No | 200 | 37.3 | 19500 | 14.251 | 51.887 |
| Catarrhini | *Papio hamadryas* | No | 197 | 37 | 12671 | 13.629 | 46.276 |
| Catarrhini | *Gorilla gorilla* | No | 202 | 35.5 | 139842 | 17.093 | 77.524 |
| Catarrhini | *Homo sapiens* | No | 208 | 37 | 70000 | 16.095 | 68.522 |
| Catarrhini | *Pan troglodytes* | No | 206 | 35.7 | 44984 | 15.457 | 62.766 |
| Catarrhini | *Pongo abelii* | No | 204 | | 60000 | 15.873 | 66.519 |
| Platyrrhini | *Alouatta palliata* | No | 174 | 36 | 4670 | 12.189 | 76.957 |
| Platyrrhini | *Aotus nancymaae** | No | 177 | | 788 | 9.622 | 49.173 |
| Platyrrhini | *Aotus trivirgatus** | No | 177 | 38 | 914.5 | 9.837 | 51.497 |
| Platyrrhini | *Callithrix jacchus* | No | 172 | 36 | 190 | 7.570 | 26.960 |
| Platyrrhini | *Saimiri bolivensis** | No | 174 | | 615 | 9.264 | 45.301 |
| Platyrrhini | *Saimiri sciureus** | No | 174 | 38 | 836.7 | 9.709 | 50.109 |
| Lemuriformes | *Cheirogaleus medius* | Yes | 166 | 38 | 300 | 8.229 | 39.027 |
| Lemuriformes | *Eulemur fulvus* | No | 180 | 36.5 | 2374 | 11.213 | 60.483 |
| Lemuriformes | *Eulemur mongoz* | No | 180 | | 1350 | 10.399 | 54.630 |
| Lemuriformes | *Indri indri* | No | 180 | | 7750 | 12.920 | 72.756 |
| Lemuriformes | *Lemur catta* | No | 178 | | 2200 | 11.103 | 59.692 |
| Lemuriformes | *Microcebus griseorufus* | Yes | 174 | | 62.5 | 5.966 | 22.756 |
| Lemuriformes | *Microcebus murinus* | Yes | 174 | 35.7 | 62.5 | 5.966 | 22.756 |
| Lemuriformes | *Microcebus ravelobensis* | Yes | 173 | | 71.5 | 6.160 | 24.152 |
| Lemuriformes | *Mirza coquereli* | Yes | 176 | | 310 | 8.276 | 39.367 |
| Lemuriformes | *Prolemur simus* | No | 180 | | 2350 | 11.198 | 60.375 |
| Lemuriformes | *Propithecus coquereli** | No | 180 | | 4000 | 11.966 | 65.897 |
| Lemuriformes | *Propithecus verreauxi** | No | 180 | 36 | 3350 | 11.710 | 64.056 |
| Lemuriformes | *Varecia variegata* | No | 181 | | 3350 | 11.710 | 64.056 |
| Lorisiformes | *Arctocebus calabarensis* | No | | 36 | 206 | 7.687 | 28.393 |
| Lorisiformes | *Galago moholi** | Yes | 165 | | 170 | 7.409 | 23.982 |
| Lorisiformes | *Galago senegalensis** | No | 165 | 37.9 | 171.5 | 7.422 | 24.184 |
| Lorisiformes | *Loris tardigradus* | Yes | 166 | 35.5 | 248 | 7.954 | 32.654 |
| Lorisiformes | *Nycticebus bengali* | No | 164 | | 1500 | 10.551 | 73.987 |
| Lorisiformes | *Nycticebus coucang* | No | 167 | 35.4 | 1128.6 | 10.140 | 67.445 |
| Lorisiformes | *Otolemur crassicaudatus** | No | 166 | 36.6 | 993.5 | 9.956 | 64.523 |
| Lorisiformes | *Otolemur garnettii** | No | 166 | 36 | 1314 | 10.360 | 70.947 |
| Lorisiformes | *Perodicticus potto* | No | | 36.1 | 968.6 | 9.920 | 63.940 |

**Table S9.** Comparison between Chickens and Mammals of Metabolic Rates and Rates of Protein Synthesis. Data for laying hens and mammals (Muramatsu, 1990)[21] were corrected for egg production: -14% of the total rate of protein synthesis, and -3% of *bMR* (Hiramoto et al, 1989[22] and Sakomura, 2004[23], respectively).

| ***Species*** | ***Mass, kg*** | ***Specific Rate of Protein Synthesis g/kg$^{0.75}$/day*** | ***Ratio*** | ***Species*** | ***Mass, kg*** | ***rMR ml O2/h*** | ***Specific Metabolic Rate ml $O_2$/kg$^{0.75}$/h*** | ***Ratio*** |
|---|---|---|---|---|---|---|---|---|
| Chicken (avg) | 1.49 | 26.00 | Chicken/ Mammal | Chicken | 1.493 | 1,219 | 5.07 | Chicken/ Mammal |
| Mouse | 0.03 | 18.60 | 1.40 | Mouse | 0.03 | 310.0 | 3.85 | 1.32 |
| Rat | 0.51 | 17.40 | 1.49 | Rat | 0.25 | 309.1 | 3.82 | 1.33 |
| Rabbit | 3.60 | 16.40 | 1.59 | Rabbit | 2.15 | 312.0 | 4.08 | 1.17 |
| Human (avg) | 66.5 | 14.60 | 1.78 | Human | 60.50 | 311.6 | 3.52 | 1.44 |
| Cow (avg) | 564 | 16.10 | 1.61 | Cow | 347.00 | 53,442 | 3.74 | 1.36 |

**Table S10**. Markers of Constraint and Adaptation in Mammals with High- versus Low- log $C_{min}$ Residuals. Data represent mean values (s.d.) and significance of differences by analysis of variance. Values of $U_{CT}$ and *cMR.71* were drawn from Khaliq et al (2014)[24] while values for $T_b$ and $T_a$ were drawn from Clarke et al (2010)[18]. These data provide statistical support for the interpretation of main text Figure 11.

| ***Group*** | ***N*** | ***$U_{CT}$*** | ***P*** | ***cMR.71*** | ***P*** | ***$T_b$*** | ***P*** | ***$T_a$*** | ***P*** |
|---|---|---|---|---|---|---|---|---|---|
| High Cmin Res. | 34 | 34.27 (2.92) | **3.54e-07** | 0.090 (0.040) | **11.36e-04** | | | | |
| Low Cmin Res. | 239 | 31.56 (2.82) | | 0.070 (0.025) | | | | | |
| High Cmin Res. | 34 | | | | | 37.11 (1.48) | **8.78e-03** | 15.76 (8.08) | **0.4262** |
| Low Cmin Res. | 166 | | | | | 36.36 (1.51) | | 14.65 (7.29) | |

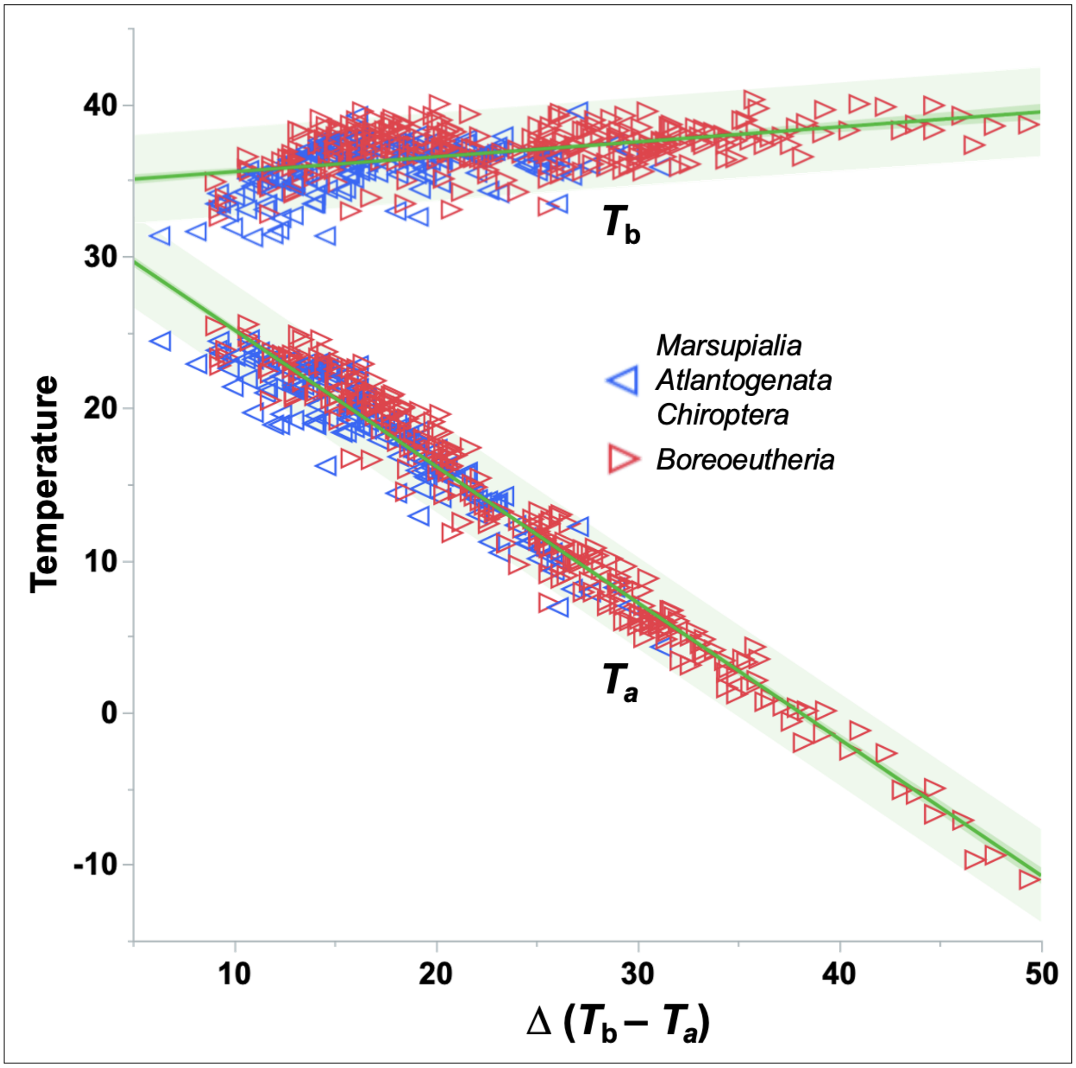

**Figure S7.** The temperature gradient that mammals can maintain varies inversely with ambient temperature, not body temperature.
Data (n = 462, excluding 52 outliers) were drawn from Clarke et al (2010)[18]. Boreoeutheria exclude Chiroptera. These data inform the interpretation of main text Figures 9 - 11.

**REFERENCES** (Supplementary Information)

17. McGeary SE, Lin KS, Shi CY, Pham TM, Bisaria N, Kelley GM and Bartel DP. 2019. The biochemical basis of microRNA targeting efficacy. *Science* 366: eaav1741.

18. Clarke A, Rothery P and Isaac NJP. 2010. Scaling of basal metabolic rate with body mass and temperature in mammals. *J Animal Eco*l 79: 610-619.

19. Pagel M, Meade A and Barker D. 2004. Bayesian estimation of ancestral character states on phylogenies. *Systematic Biol* 53: 673-684.

20. Umu SU, Paynter VM, Trondsøn H, Buschmann T, Rounge TB, Peterson KJ and Fromm B. 2023. Accurate microRNA annotation of animal genomes using trained covariance models of curated microRNA complements in MirMachine. *Cell Genomics* 3, 100348.

21. Muramatsu T. 1990. Nutrition and whole-body protein turnover in the chicken in relation to mammalian species. Nutr Res Rev 3: 211-228.

22. Hiramoto K, Muramatsu T and Okumura JI. 1989. Protein synthesis in several tissues of laying hens. *Jpn Poul Sci* 26: 340-347.

23. Sakomura NK. 2004. Modeling energy utilization in broiler breeders, laying hens and broilers. *Brazil J Poultry Sci* 6: 1-11.

24. Khaliq I, Hof C, Prinzinger R, Böhning-Gaese K and Pfenninger M. 2014. Global variation in thermal tolerances and vulnerability of endotherms to climate change. *Proc Roy Soc B* 281: 20141097 (1-8).
